# The increase of Cyclin A/cdk activity and of FAM122A-dependent inhibition of PP2A-B55 are the key events triggering mitosis

**DOI:** 10.1101/2023.06.20.545672

**Authors:** Benjamin Lacroix, Suzanne Vigneron, Jean Claude Labbé, Lionel Pintard, Gilles Labesse, Anna Castro, Thierry Lorca

**Author notes:** ^&^Both authors contributed equally to this work (Benjamin Lacroix, Suzanne Vigneron).

## Abstract

Entry into mitosis has been classically attributed to the activation of cyclin B/cdk1 amplification loop by a partial pool of this kinase that becomes active at the end of G2. However, how this pool is activated is still unknown. Here we discovered a new role of the recently identified PP2A-B55 inhibitor FAM122A in triggering mitotic entry. Accordingly, the depletion of the orthologue of FAM122A in *C. elegans*, prevents entry into mitosis in germline stem cells. Moreover, our data in Xenopus egg extract strongly supports that FAM122A-dependent inhibition of PP2A-B55 could be the initial event promoting mitotic entry. The inhibition of this phosphatase allows the subsequent phosphorylation of first mitotic substrates by cyclin A/cdk resulting in cyclin B/cdk1 and Greatwall (Gwl) activation. However, interestingly, from Gwl activation, Arpp19/ENSA become phosphorylated and compete with FAM122A promoting its dissociation from PP2A-B55 and taking over its inhibition until the end of mitosis.

## INRODUCTION

Mitotic entry and progression are induced by massive protein phosphorylation resulting from the fine-tuned balance between the kinase cyclin B/cdk1 and its counteracting phosphatase PP2A-B55^1–4^. Cyclin B/cdk1 activity is maintained low during G2 by the Wee1/Myt1 kinases that phosphorylate cdk1 on its inhibitory site tyrosine 15. At M phase entry, cyclin B/cdk1 activity is triggered by a positive feed-back loop^5, 6^. A partial active pool of this kinase phosphorylates Wee1/Myt1, as well as the Cdc25 phosphatase responsible of cdk1-tyrosine 15 dephosphorylation, promoting a rapid and complete activation of the cyclin B/cdk1 complex. How cyclin B/cdk1 partial activation is firstly triggered at G2-M is a main question yet to be answered. Inhibition of PP2A-B55 was proposed as the putative cause inducing partial cyclin B/cdk1 pool activation and mitotic entry. PP2A-B55 is regulated by the Greatwall (Gwl)-Arpp19/ENSA pathway^1, 2, 7, 8^. During G2, the activity of this phosphatase is high. Then, at mitotic entry, Gwl is activated and phosphorylates its substrates Arpp19/ENSA converting them into high affinity inhibitors of PP2A-B55. Decreased PP2A-B55 activity could then favour a partial Wee1/Myt1/Cdc25 phosphorylation and reactivation of cyclin B/cdk1 triggering the feedback loop. However, Gwl activation itself depends on cyclin B/cdk1 ruling out the modulation of this phosphatase as the first event triggering cyclin B/cdk1 firing and mitotic entry^9, 10^. Moreover, although both Gwl and Arpp19/ENSA are essential for mitotic entry in Xenopus egg extract model^2, 4^, the depletion of these proteins do not prevent G2-M transition in human and mouse cells^8, 11^.

Besides cyclin B/cdk1, cyclin A/cdk is also present and is a poor substrate of Myt1 kinase during G2^12, 13^. This kinase could participate to Wee1/Myt1/Cdc25 phosphorylation during G2-M if the activity of PP2A-B55 is partially inhibit during this transition. In this line, FAM122A has been recently described as a new inhibitor of PP2A-B55 that is present and active during G2.

FAM122A was first identified as an interactor of the PP2A-B55 complex that inhibits this phosphatase and promotes the degradation of PP2A catalytic subunit.^14^ This protein was additionally shown to negatively modulate PP2A-B55α in the nucleus. Its phosphorylation by Chk1 and its subsequent retention into the cytoplasm is essential for the G2/M checkpoint by preventing the nuclear inhibition of this phosphatase and permitting the dephosphorylation and stabilisation of Wee1^15^. By its presence in G2 and its capacity to specifically inhibit PP2A-B55, FAM122A is a good candidate to fulfil the role of PP2A-B55 inhibitor able to potentiate cyclin A/cdk-dependent phosphorylation at G2 and to trigger M phase. We thus investigated the role of this protein in mitotic entry. Using Xenopus egg extracts we demonstrate that the inhibition of PP2A-B55 by FAM122A does not induce Wee1 or PP2A catalytic subunit proteolysis. Conversely, it promotes mitotic entry when added to interphase extracts. FAM122A does not require its phosphorylation or the activity of Gwl or cyclin B/cdk1 to induce first mitotic substrate phosphorylation, however, it depends on cyclin A/cdk activity, indicating that FAM122A promotes entry into mitosis via this cyclin/cdk complex. Upon addition to interphase egg extracts, FAM122A first binds and inhibits PP2A-B55 and promotes cyclin A/cdk1 dependent phosphorylation of mitotic substrates and the final activation of cyclin B/cdk1 and Gwl. Interestingly, once Gwl is active and phosphorylates Arpp19, this inhibitor competes with FAM122A for its binding to PP2A-B55 and promotes its dissociation. By comparing predicted structure obtained by AlphaFold v2.2, we could predict that FAM122A inhibits the phosphatase by blocking substrate access to the catalytic site while phospho-Serine Gwl site of Arpp19 makes stronger and deeper interactions with manganese ions of the catalytic site thus explaining why it is able to displace FAM122A by competition. The role of FAM122A on mitosis appears to be evolutionarily conserved over species since its knockdown prevents normal G2-M transition and mitotic progression in germline stem cells of *C. elegans*. Thus, our data suggest that cyclin A-Cdk1 activation and FAM122A accumulation during G2 could be the first events triggering mitotic entry.

## RESULTS

### PP2A-B55 inhibition by FAM122A does not involve changes in PP2A-C or Wee1 levels

The role of FAM122A as a specific PP2A-B55 inhibitor has been reported by two different laboratories^14, 15^. One laboratory reported a role of this protein in the negative modulation of PP2A-B55 via the degradation of the PP2A-B55 catalytic subunit C^14^. The second one suggested that the inhibition of PP2A-B55 by FAM122A induces the dephosphorylation and degradation of Wee1. We checked these two premises in interphase Xenopus egg extracts (interphase extracts), in which protein phosphorylation mimics the one observed in G2 cells. We, therefore, added either Xenopus or human histidine recombinant FAM122A proteins to these extracts and measured the levels of PP2A-C and Wee1 over time. Surprisingly, we could not observe any variation of the amount of these two proteins in the extracts (Figure 1a). We next proceeded by checking whether our recombinant FAM122A forms can inhibit PP2A-B55. To do this, we used interphase egg extracts in which ATP was eliminated and thus endogenous kinases are inactive (called hereafter as kinase-inactivated extracts). Because of the absence of endogenous active kinases, substrate dephosphorylation in these extracts directly reflects phosphatase activity(ies). Kinase-inactivated extracts were supplemented, or not, with the corresponding FAM122A recombinant protein (human/Xenopus) and subsequently with recombinant Arpp19 pre-phosphorylated at S113 by PKA known to be a bona fide PP2A-B55 substrate^16, 17^. Dephosphorylation of S113 of Arpp19 was then followed by western blot over time. As expected by its capacity to inhibit PP2A-B55, both human and Xenopus FAM122A drastically delayed Arpp19 dephosphorylation (Figure 1b). Similar results were observed using phosphorylated-T481 PRC1, another well-established substrate of PP2A-B55 phosphatase^18^ was used. We next checked the association of His-FAM122A to PP2A-B55 by using histidine-pull down. B55, A and C subunits of PP2A were present in the histidine but not in the control pull-down of Xenopus and human FAM122A (Figure 1c). Thus, FAM122A can directly bind and inhibit PP2A-B55 without any modification of the levels of the catalytic subunit of this phosphatase.

**Figure 1.**
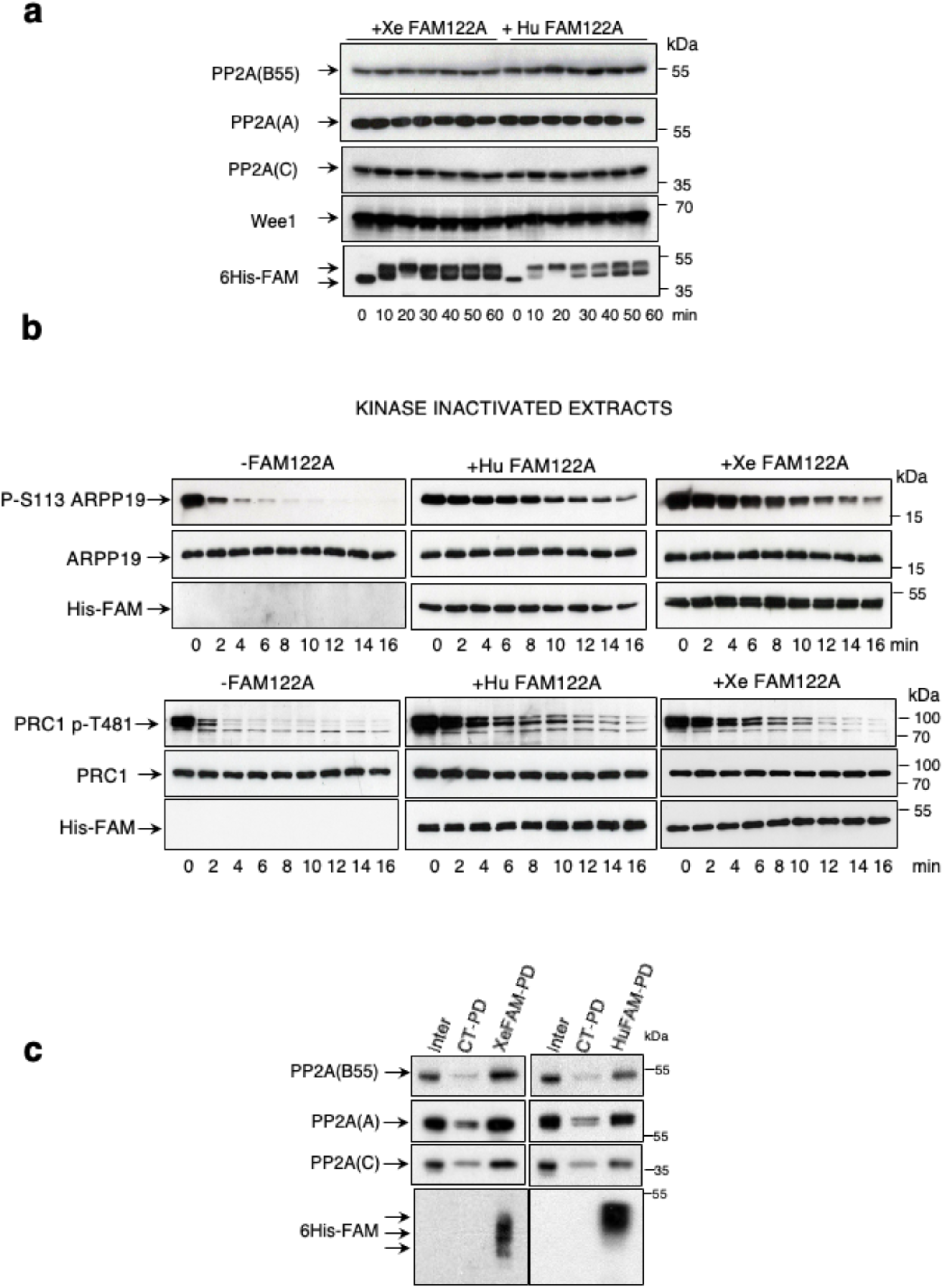
PP2A-B55 inhibition by FAM122A does not involve changes in PP2A-C or Wee1 levels. **(a)** 20 μl of interphase extracts were supplemented with 2 μl (1,6 μg) of either Xenopus or human FAM122A and 0,8 μl of the mix was recovered at different time points to measure the indicated proteins by western blot. **(b)** Arpp19 or PRC1 proteins *"in vitro"* phosphorylated by PKA or purified cyclin A/cdk1 respectively were supplemented together with Xenopus or human FAM122A to kinase inactivated interphase extracts and the dephosphorylation rate of S113 of Arpp19 and T481 of PRC1 as well as the levels of Xenopus and human FAM122A were analysed by western blot. **(c)** Interphase extracts were supplemented (Xe/HuFAM-PD) or not (CT-PD) with either Xenopus or human FAM122A as in (a) and used for histidine pull-down. The levels of PP2A B55, A and C subunits as well as the amount of FAM122A were checked by western blot.

### FAM122A triggers mitotic entry by inhibiting PP2A-B55 and permitting cyclin A/cdk-dependent phosphorylation of mitotic substrates

We next checked the impact of FAM122A on mitotic progression by measuring the phosphorylation of mitotic substrates upon the addition of this protein into interphase extracts. FAM122A induced a first phosphorylation of Gwl, the Anaphase Promoting Complex (APC) subunit Cdc27, and of Cdc25 concomitantly with the activation of cyclin B/cdk1 as reflected by the loss of the phosphate on tyrosine 15 inhibitory site of cdk1. This was followed by the subsequent degradation of cyclin A and B and the dephosphorylation of the above indicated substrates revealing that FAM122A promotes mitotic entry but is unable to maintain the mitotic state (Figure 2a). We then sought to assess whether, as for Arpp19, FAM122A requires phosphorylation by Gwl to stably bind and inhibit PP2A-B55. Against this hypothesis, our data indicate that Gwl is unable to phosphorylate FAM122A "*in vitro*" (Figure 2b) and the depletion of this kinase in interphase extracts did not prevent mitotic entry induced by FAM122A (Figure 2c). Moreover, a FAM122A form in which all serine and threonine residues were mutated into alanine kept its capacity to induce mitosis (Figure 2d) and displayed a similar PP2A-B55 binding capacity than the wildtype FAM122A (Figure 2e). These data fully rule out a role of its phosphorylation in PP2A-B55 binding and inhibition. Thus, FAM122A requires neither phosphorylation nor Gwl activity to trigger mitotic entry.

**Figure 2.**
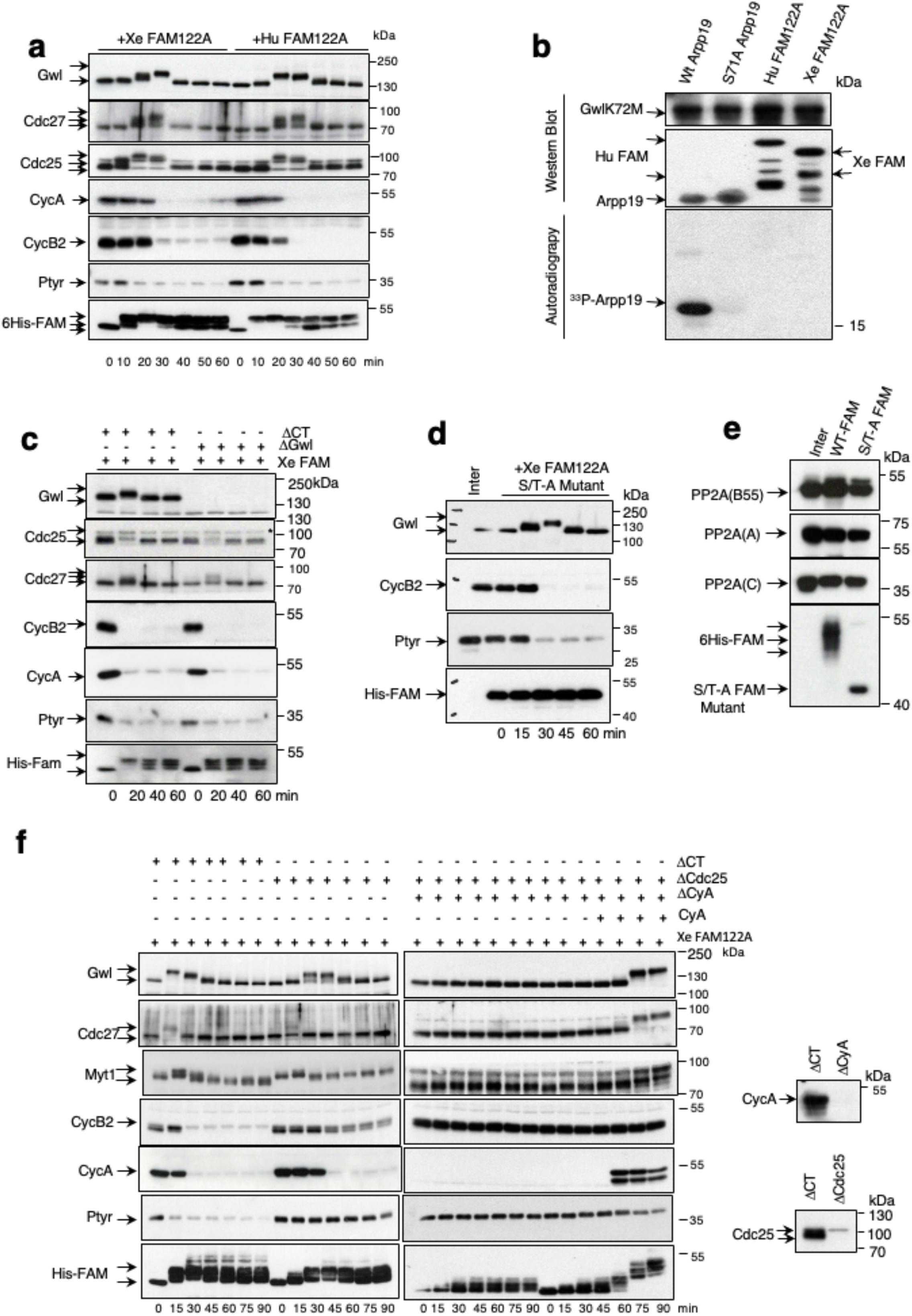
FAM122A triggers mitotic entry by inhibiting PP2A-B55 and permitting cyclin A/cdk-dependent phosphorylation of mitotic substrates. **(a)** Extracts were treated as for Figure 1a and the levels of the indicated proteins as well as the phosphorylation of cdk1 on tyrosine 15 were checked by western blot. **(b)** His-wildtype Arpp19, the Gwl phosphorylation site mutant (Serine 71 -to-Alanine) of this protein, His-human FAM122A and His-Xenopus FAM122A were phosphorylated "*in vitro*" by GST-human GwlK72M hyperactive mutant in a final phosphorylation reaction mixture of 10 μl as indicated in Methods. 5 μl were then used for western blot using anti-histidine antibodies to detect Arpp19 and human and Xenopus FAM122A levels or anti-Gwl antibodies to detect Gwl amount. The rest was used to detect γ^33^P by autoradiography. **(c)** Interphase extracts were depleted with a control or anti-Gwl antibodies and supplemented with Xe FAM122A. Samples were then analysed over time for the levels and phosphorylation of the indicated proteins. **(d)** Interphase extracts were supplemented with the Xenopus FAM122A protein in which all serine/threonine residues were mutated into alanine. Samples were recovered at the indicated time points and used for western blot. **(e)** Interphase extracts supplemented with a wildtype or a Xenopus FAM122A protein in which all serine/threonine residues were mutated into alanine and used for histidine pulldown and western blot to measure their association with B55, A and C subunits of PP2A. **(f)** Interphase extracts were depleted using control or anti-Cdc25 antibodies, or s subjected to two consecutive depletion using anti-Cdc25 and cyclin A. Depleted supernatants were then supplemented with Xe FAM122A protein. When indicated, 180 ng of purified cyclin A was added to the mix. Samples were recovered over time and used for western blot. Depletion of cyclin A and Cdc25 in the extracts was confirmed by western blot.

We then investigated whether mitotic substrate phosphorylation induced by FAM122A requires cyclin B/cdk1 activation by examining the effect of depleting Cdc25, the phosphatase dephosphorylating tyrosine 15 of cdk1 and triggering cyclin B/cdk1 activity. As expected, Cdc25-devoid extracts displayed inactive cyclin B/cdk1, as stated by the presence of the phospho-tyrosine 15 on cdk1. Moreover, cyclin A became fully proteolyzed whereas cyclin B remained mostly stable although we detected a loss of a small fraction of this protein likely due to the phosphorylation, under these conditions, of Cdc27 and the activation of the APC (Figure 2f). Besides Cdc27, FAM122A, via PP2A-B55 inhibition, was also able to promote the phosphorylation and activation of Gwl and, most importantly, the phosphorylation and inactivation of the cdk1 inhibitory kinase Myt1. Since due to Cdc25 depletion cyclin B/cdk1 was inactive in these extracts, we attributed these phosphorylations to cyclin A/cdk1. In this line, we previously showed that this complex escapes to Wee1-Myt1 inhibitory phosphorylation and is fully active in interphase extracts^19^. To confirm this hypothesis, we co-depleted Cdc25 and cyclin A in FAM122A-suplemented interphase extracts. Remarkably, the loss of these two proteins prevented substrate phosphorylation resulting from FAM122A-dependent inhibition of PP2A-B55 (Figure 2f). However, this phosphorylation was restored again if 45 minutes upon Cdc25/cyclin A co-depletion and FAM122A addition, cyclin A was supplemented again to these extracts. These data demonstrate that the decrease of the PP2A-B55 activity in interphase extracts induced by FAM122A is sufficient to permit cyclin A-cdk1-dependent phosphorylation that will then trigger cyclin B/cdk1 reactivation. Interestingly, this suggests that an increase of the endogenous levels of either FAM122A or cyclin A would be sufficient to trigger mitotic entry and would thus escape to the negative regulation of active PP2A-B55 during interphase.

### Molecular determinants of FAM122A controlling the association and the inhibition of PP2A-B55

AlphaFold artificial intelligence software (Alphabet’s/Google’s DeepMind) predicts the presence of two αhelices in the FAM122A protein, α-helix1: residues 84-93 (human)/71-82 (Xenopus) and α-helix2: residues 96-119 (human)/85-110 (Xenopus) (Figure 3a). In order to investigate if these structured regions could participate to the association of this protein to PP2A-B55, we constructed a FAM122 Xenopus mutant forms deleted of either α-helix1 (αH1) or α-helix2 (αH2) and we measured their capacity to induce mitotic entry and to bind to PP2A B55 and C subunits when added to interphase extracts. Neither of these two mutants were able to induce mitosis (Figure 3b) or to bind to PP2A-B55 (Figure 3c) indicating that these two regions are essential for FAM122A inhibitory activity. We next constructed a N-terminal Δ(1-72) and a C-terminal (Δ110-270) mutant forms in order to determine whether any of these regions could additionally participate to the functionality of this protein. Unlike the two α-helix regions, these two mutants displayed similar properties than the wildtype form (Figure 3d and e). Finally, we proceed to the construction of FAM122A single mutant forms for all conserved residues present in αH1 and αH2 regions. Data on mitotic entry and PP2A-B55 binding capacities of these mutants are shown in Supplementary Figure 1a and 1b and summarized in Figure 3f. Residues of αH1, R73, L74, I77 and E80 were essential for FAM122A inhibitory activity. Indeed, αH1 sequence SRLHQIKQEE, is close to the previous reported Short Linear Motif (SLiM) designed for PP2A-B55, with consensus sequence p[ST]-P-x(4,10)-[RK]-V-xx-[VI]-R^20^ and is highly conserved between species (Supplementary Figure 1c). This motif is present in PP2A-B55 substrates and is required for their binding to the phosphatase via their interaction with the B55 subunit. As so, these residues are involved in FAM122A association to B55. For αH2 region, we identified positions E90, H93, E94 and R95 as being essential for FAM122A activity. Thus, we pinpointed these two structured regions of FAM122A essential for its function and we identified the key residues directly mediating PP2A-B55 phosphatase interaction.

**Figure 3.**
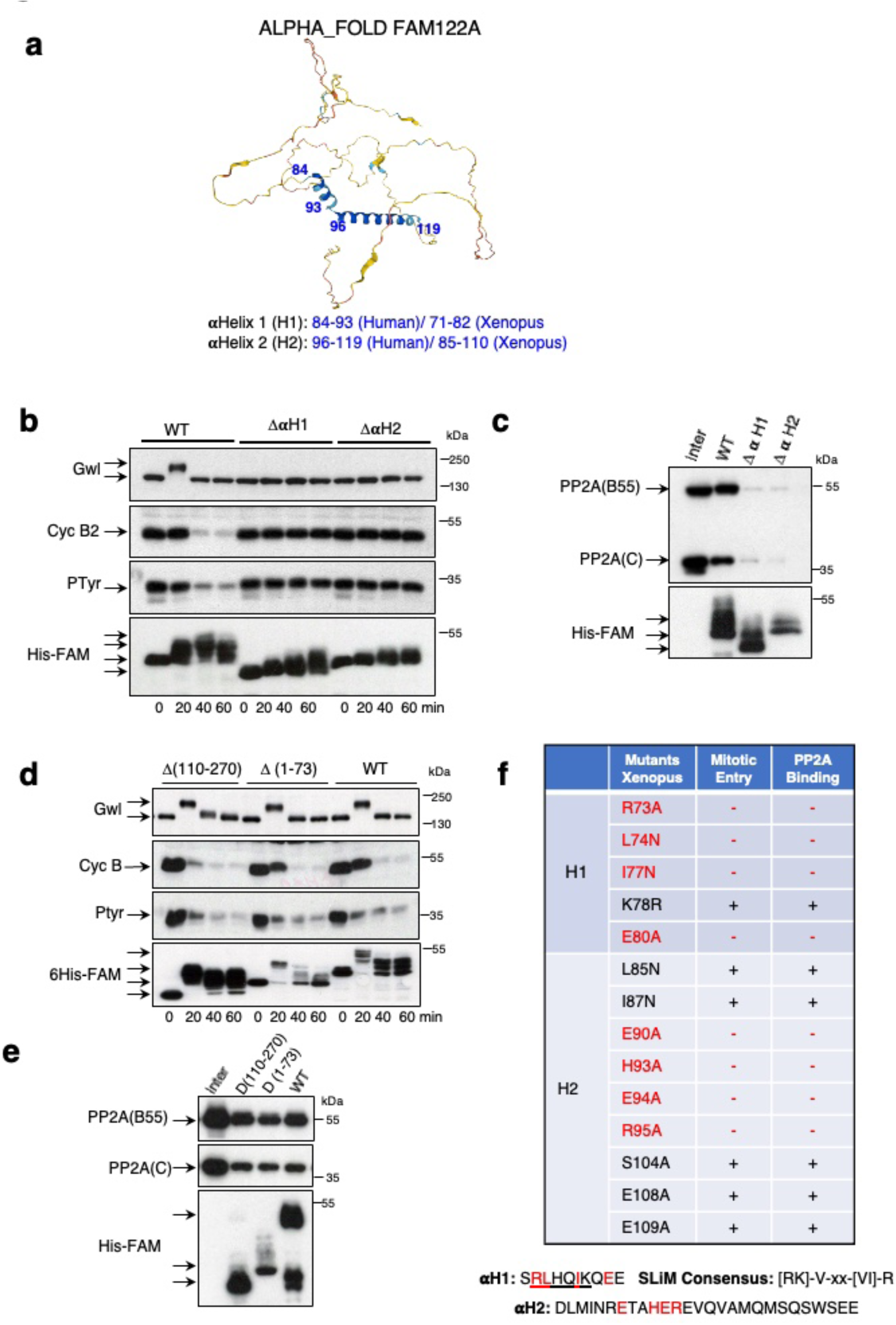
Molecular determinants controlling FAM122A inhibitory activity. **(a)** AlphaFold Monomer v2.0 predicts two structured αhelices (αHelix 1 and αHelix 2) in the human FAM122A protein **(b)** Interphase extracts were supplemented with the wildtype or the αH1 or αH2 mutants of Xe FAM122A and the phosphorylation and levels of the indicated proteins measured at the indicated times. **(c)** Wildtype and αH1 or αH2 mutants supplemented interphase extracts were used for His-pulldown to measure the association of these proteins to B55 and C subunits of PP2A-B55. **(d)** The wildtype and the indicated mutant forms of Xe FAM122A were supplemented to interphase extracts and used for western blot. **(e)** His-pulldown was performed in extracts treated as in (d) to check the binding of the different forms of Xe FAM122A form to B55 and C. **(f)** Table indicating the capacity of the different single mutants of Xe FAM122A to induce mitotic entry and to bind to PP2A-B55. Mutants that have lost their inhibitory activity are depicted in red. The sequence of the αH1, the SLiM consensus sequence and the sequence of αH2 are shown. Residues essential to keep the inhibitory activity of Xe FAM122A are highlighted in red.

### Gwl activation at mitotic entry promotes the dissociation of FAM122A from PP2A-B55

Data above demonstrate that the addition of FAM122A to interphase extracts promotes mitotic entry but is unable to maintain mitotic substrate phosphorylation. Since the levels of FAM122A remain constant throughout the experiment and that FAM122A activity appears not to be impacted by phosphorylation, we sought to understand why FAM122A is not able to maintain the mitotic state. To do so, we first checked the capacity of Xenopus and human FAM122A proteins to bind PP2A-B55 when they were supplemented to metaphase II arrested extracts (CSF extracts) which mimics the mitotic state. Interestingly, B55 and C subunit binding to FAM122A is clearly observed during interphase, whereas no association for B55 and a drastic decrease for C catalytic subunit were observed in CSF extracts (Figure 4a).

**Figure 4.**
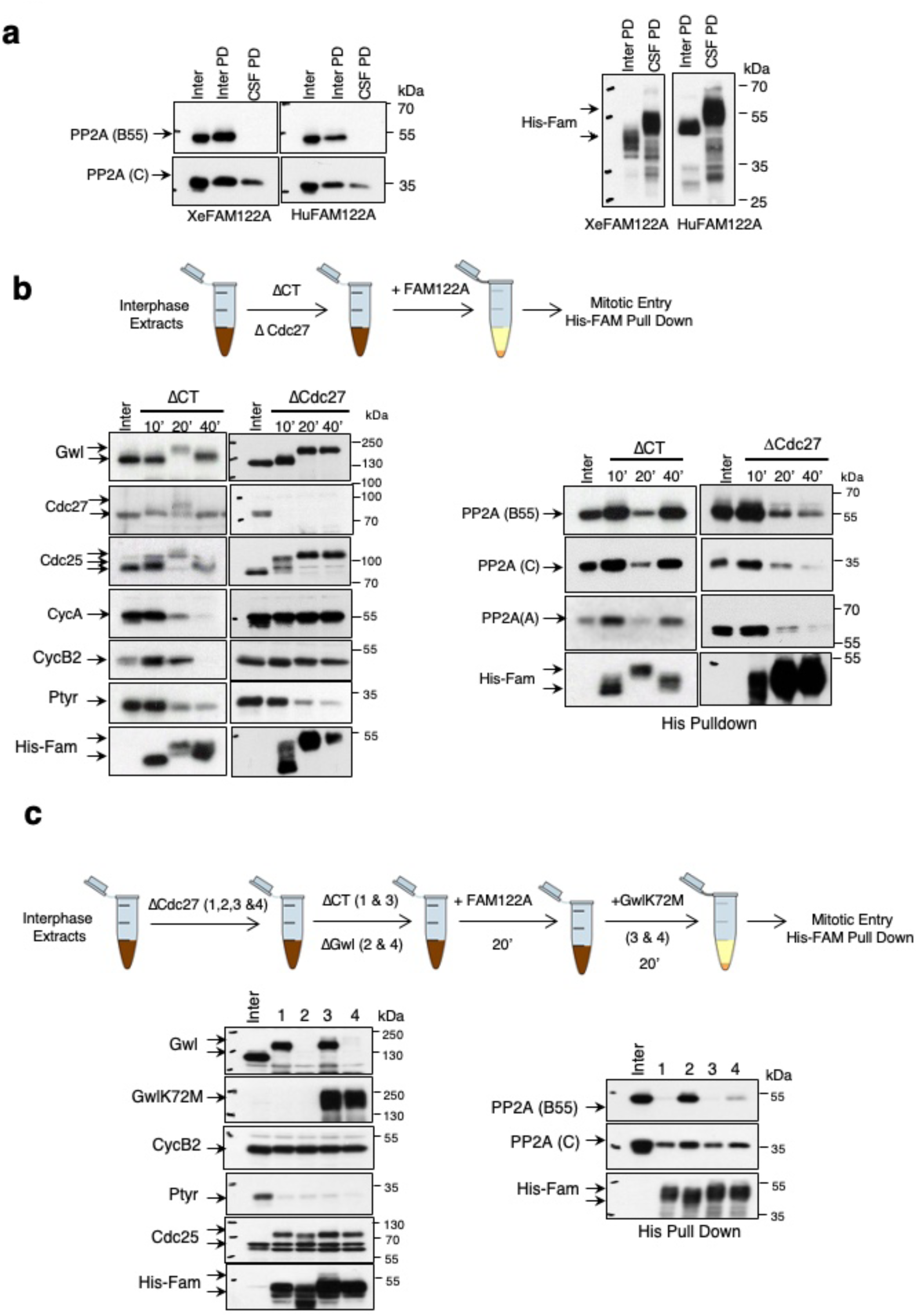
Gwl activation at mitotic entry promotes the dissociation of FAM122A from PP2A-B55. **(a)** Interphase and CSF extracts were supplemented with Xe FAM122A and Hu FAM122A and used for His-pulldown to measure the association of this protein with B55 and C subunits of PP2A-B55. The levels of His-Xe/HuFAM122A recovered in the pull down are also shown. Inter PD: Interphase Pull Down. CSF PD: CSF pull down. **(b)** Interphase extracts were depleted with control or with Cdc27 antibodies and subsequently supplemented with Xe FAM122A and used for western blot or for His-pulldown to assess the binding of Xe FAM122A to PP2A-B55. **(c)** Interphase extracts were depleted of Cdc27, subjected to a second immunodepletion using control (lanes 1 & 3) or anti-Gwl antibodies (lanes 2 & 4) and supplemented with Xe FAM122A. 20 minutes later Gwl activity was restored (lanes 3 & 4) or not (lanes 1 & 2) by adding a hyperactive form of Gwl (GwlK72M). Samples were used for western blot and His-pulldown.

To corroborate these observations, we tested whether FAM122A would be dissociated from B55 subunit in extracts blocked in mitosis. In order to obtain mitotic blocked extracts, we prevented cyclin B degradation in interphase extracts by the depletion of the APC subunit Cdc27 and we then added FAM122A. FAM122A addition promoted mitotic entry but, because of the incapacity to proteolyze cyclin B, extracts remained blocked in this phase of the cell cycle. Under these conditions, we measured the temporal pattern of protein phosphorylation and association of FAM122A to PP2A-B55. As expected, addition of FAM122A to control interphase extracts promoted mitotic entry and exit, while mitosis was maintained throughout the experiment when Cdc27 was depleted (Figure 4b). In control extracts, FAM122A rapidly associated to B55, A and C subunits from its addition, then dissociated concomitantly with mitotic entry and re-associated again upon exit of mitosis by cyclin B degradation (Figure 4b). Conversely, in Cdc27-depleted interphase extracts, FAM122A also displayed a first association to these proteins but then, definitively lost this interaction from the establishment of the mitotic state. Taken together, these observations suggest that FAM122A-B55 binding is prevented during mitosis.

Besides, cyclin B/cdk1, the Gwl kinase is also required to maintain the mitotic state. Although our data shown above demonstrate that Gwl cannot phosphorylate FAM122A to promote mitotic entry, we tested whether the PP2A-B55 inhibitory activity of FAM122A could be indirectly modulated by this kinase once the mitotic state is achieved. To this, we repeated the previous experiment except that extracts were immunodepleted of both, Cdc27 and the Gwl kinase. As expected by the depletion of Cdc27, all extracts entered into mitosis and kept the mitotic state as shown by the stability of cyclin B, the continuous phosphorylation of Cdc25 and the loss of phospho-tyrosine 15 cdk1 signal throughout the experiment (Figure 4c). Interestingly, unlike Cdc27/control-co-depleted extracts (Figure 4c, lane 1, His Pull Down) and despite the presence of a fully active cyclin B/cdk1 complex, FAM122A was able to bind to B55 and to increase its association to PP2A C subunit in Cdc27/Gwl-co-depleted extracts (Figure 4c, lane 2, His Pull Down). However, FAM122A dissociated again from B55 when a Gwl hyperactive form was further supplemented (Figure 4c, lane 4, His Pull Down). Thus, FAM122A dissociation from PP2A during mitosis is modulated by the Gwl kinase.

### Arpp19 competes and dissociates FAM122A from PP2A-B55 during mitosis

We next sought to decipher how Gwl could regulate FAM122A association to PP2A-B55. Gwl phosphorylates Arpp19 during mitosis, turning it into a high affinity interactor of PP2A-B55^1, 2, 21^. We thus, asked whether during mitosis, phosphorylated Arpp19 competes with FAM122A for PP2A-B55 binding. To test this hypothesis, we supplemented interphase extracts with FAM122A and 40 minutes later, once the extract returned to an interphase state and FAM122A binds again PP2A-B55 (Figure 2a and 4b), it was either (1) used to perform a histidine-FAM122A pulldown and subsequently supplemented with thio-S71-phosphorylated Arpp19 or the other way around, (2) first supplemented with a thio-S71-phosphorylated Arpp19 and subsequently subjected to histidine-FAM122A pulldown (Figure 5a, scheme). Finally, FAM122A association to B55 was measured. Strikingly, the presence of a thio-S71-phosphorylated form of Arpp19 (Thio-S71-Arpp19) that cannot be dephosphorylated induced the dissociation of FAM122A from PP2A-B55 in both cases confirming our hypothesis that Arpp19 displaces FAM122A from PP2A-B55 by competing for this phosphatase.

**Figure 5.**
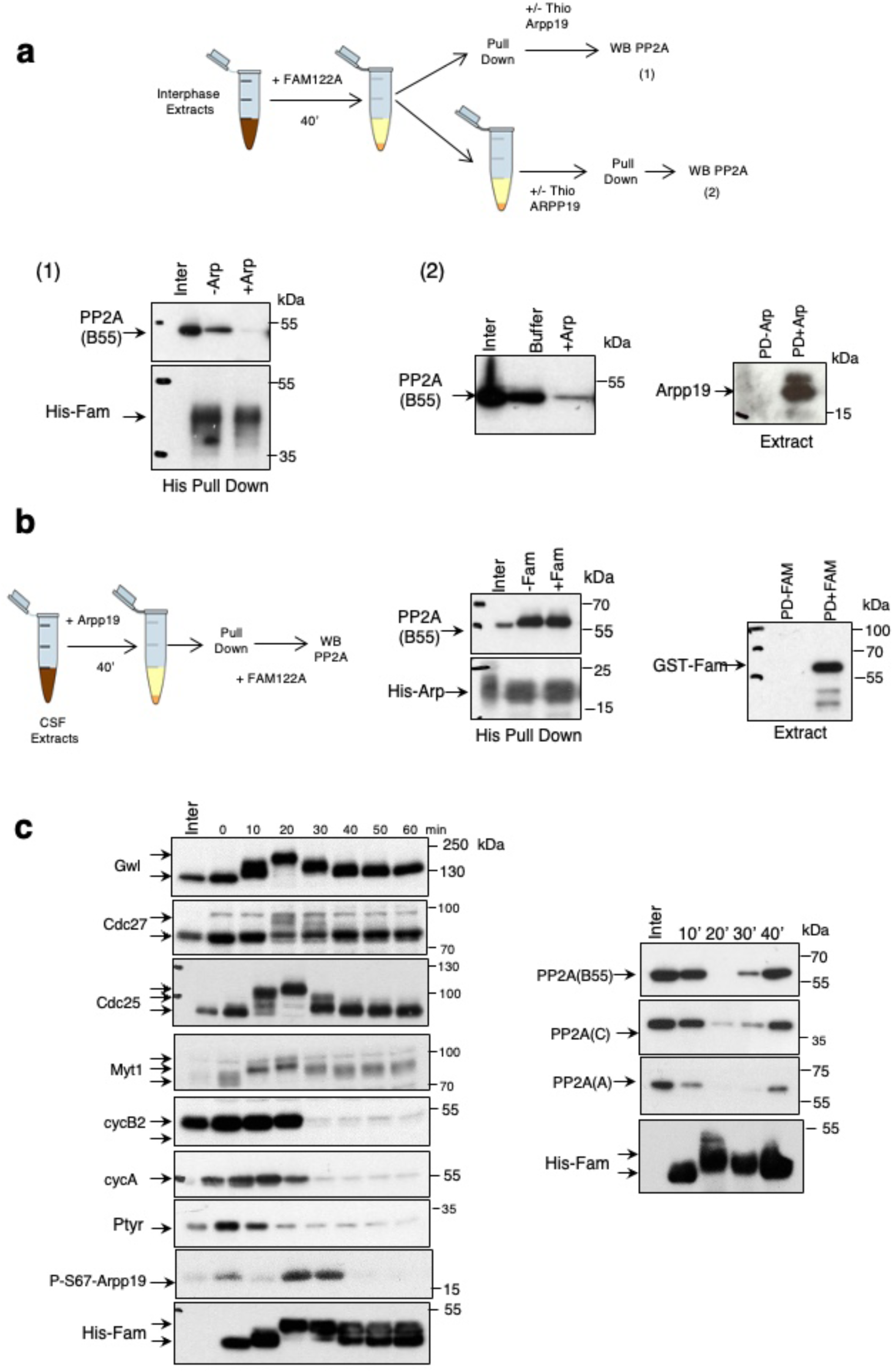
Arpp19 dissociates FAM122A from PP2A-B55 during mitosis. **(a)** Interphase extracts were supplemented with 12,87 μg of Xe FAM122A for 40 minutes and subsequently divided into two samples. One of these samples was used to perform a His-pulldown that was then supplemented with 1,04 μg of untagged Thio-Arpp19 protein and used to measure bound B55 protein. The second sample was firstly supplemented with untagged 1,04 μg of Thio-Arpp19 and subsequently used for His-pulldown to measure His-Xe FAM122A bound B55 subunit of PP2A. A sample of the His-Xe FAM122A pull downs supplemented (PD+Arp) or not (PD-Arp) with untagged Thio-Arpp19 before washing is shown. **(b)** CSF extracts were supplemented with 1,35 μg His-Arpp19 and, 40 minutes later, used for His-pulldown. Arpp19-pull downs were then supplemented with 3,6 μg GST-Xe FAM122A and the levels of B55 remaining were measured. A sample of the His-Arpp19 pull downs supplemented (PD+FAM) or not (PD-FAM) before washing is shown. **(c)** Interphase extracts were supplemented with His-Xe FAM122A and used for either western blot to measure the levels and the phosphorylation of the indicated proteins and or for His-pulldown determine Xe FAM122A binding to PP2A B55, A and C subunits at the indicated time points.

We also performed the reverse experiment and tested whether GST-FAM122A could dissociate a pArpp19 complexed to PP2A-B55 phosphatase. Thus, CSF extracts were supplemented with His-Arpp19 protein. Because of the activated Gwl kinase present in these extracts, exogenous Arpp19 is immediately phosphorylated and tightly binds PP2A-B55. pArpp19-PP2A-B55 complex was then recovered by histidine-pulldown and supplemented with high doses of GST-FAM122A. Then the level of B55 bound to pArpp19 was measured by western blot. Interestingly, the levels of B55 bound to pArpp19 did not change upon FAM122A addition indicating that FAM122A was unable to dissociate this protein from PP2A-B55 complex (Figure 5b). These data indicate that phospho-S71-Arpp19 displays a much higher affinity for PP2A-B55 than FAM122A and induces its dissociation. Supporting this hypothesis, our data above (Figure 4 and Figure 5c) demonstrate that endogenous Arpp19 is able to dissociate ectopic His-FAM122A protein from PP2A-B55 despite we estimated the molarity of the former in these experiments being 187 times lower than the latter (0.115 μM endogenous Arpp19 versus 21.5 μM ectopic His-FAM122A, Supplementary Figure 2).

Thus, Arpp19 phosphorylation during mitosis promotes the dissociation of FAM122A from PP2A-B55 and takes over the inhibition of this phosphatase. Accordingly, B55 was present in FAM122A pulldowns immediately upon being added to interphase extracts and dissociated concomitantly with Gwl and cyclin B/cdk1 activation and Arpp19 phosphorylation. This protein was re-associated again once cyclin B was degraded, Gwl was inactivated and Arpp19 was dephosphorylated at mitotic exit (Figure 5c).

### The different binding of Arpp19 and FAM122A to PP2A-B55 could explain the ability of the first inhibitor to dissociate the second from phosphatase

In order to understand how Arpp19 can dissociate FAM122A from PP2A-B55, we modeled the quaternary complex of PP2A with Arpp19 from three distant species (human, Xenopus and *C. elegans*) using Alphafold_multimer (https://www.biorxiv.org/content/early/2021/10/04/2021.10.04.463034) with default settings and no relaxation. The resulting models looks globally similar. The molecular organization of the PP2A subunits reproduces well the known crystal structure (PDB3DW8) with little unfolded regions. Arpp19 (for the human and Xenopus complexes) and the unique orthologue for Arpp19/ENSA in *C. elegans* called ENSA are mostly unfolded (Figure 6a, left panel, orange) and share three common helical stretches, labelled hereafter as α1, α2 and α3. A first helix, α0, in human Arpp19 and Xenopus Arpp19, that is not predicted in worm ENSA, seems to form no interaction with PP2A and will not be further considered. On the contrary, the three conserved helices (α1: residues 24-35, α2 :45-54 and α3 :60-73 in human Arpp19) are predicted to interact with the phosphatase. The first two contact the B55 subunit (Figure 6a, left panel, blue) while the third one points toward the catalytic site of the C subunit (Figure 6a, left panel, green). No contact is detected between the Arpp19 chains with the scaffolding subunit (Figure 6a, left panel, violet). The interaction with B55 mainly relies on the helix α1 that displays tight contacts along the helix from Arpp19. The interactions conserved among the three complexes includes two salt bridges (E25 and R35 from Arpp19 with R330 and E338 from B55 in the human complex) and hydrophobic interactions involving L32 from Arpp19 and F280, F281 and C334 from B55 subunit in human. In addition, two N-capping and one C-capping (involving E28, E29 and K26 in human and Arpp19, respectively) are predicted. The second helix of Arpp19, α2, shows more loose interactions and it seems to mainly bridge the helix α1 with the third helix. Helix α3 partially overlaps with the QKYFDSGDY motif conserved in all Arpp19 and ENSA sequences and containing the serine phosphorylated by Gwl. This phosphorylation is essential for strong inhibition of PP2A. We modeled the phosphoserine using a dedicated server (https://isptm2.cbs.cnrs.fr described in the "Methods" section) and also added the two manganese atoms within the catalytic center of the C subunit of PP2A for further analysis. Although in the human and Xenopus complexes, the serine is not predicted to directly interact with the catalytic manganese (distance PO4-Mn :∼ 8 A), such interactions appeared plausible in the worm complex (distance PO4-Mn :∼ 5 A). This direct contact would explain the strong interaction of phosphorylated Arpp19 with PP2A-B55. However, the inhibitory activity has been shown to be not only characterized by the high affinity of Arpp19 towards PP2A-B55, but also by the slow dephosphorylation of its Gwl site^16^. The present modeling does not permit to fully understand the absence of catalytic activity on the phosphorylated serine. However, this model agrees with the previously reported mutational scanning of the QKYFDSGDY motif^16^ to suggest that a precise and peculiar orientation of the phosphorylated sidechain would be required to maintain a tight but unreactive interaction. Indeed, mutations to alanine of close phosphoserine neighbor residues in this sequence dramatically accelerate Arpp19 dephosphorylation by PP2A-B55 while lowering binding at the same time^16^. Additional interactions are predicted at this interface in the worm complex (and partially reproduced in the human and Xenopus assemblies) (see Figure 6b). These interactions include two salt-bridges involving the Gwl site phosphoserine (S62) and the preceding asparate residue (D61) from human Arpp19 and two arginines of human PP2A C subunit (R89 and R214) lying at the entrance of the catalytic site. Another salt bridge may occur between another aspartate from Arpp19 (D64 in human Arpp19) and a lysine (K88) in the human B55 subunit. In addition, in the worm complex, we observed the nearby and conserved phenylalanine F65 (F60 in human Arpp19) stacked in between the subunits B and C. This residue seems to make hydrophobic contact with a leucine L126 of B55 (L87 in human B55). Other contacts are not systematically predicted in the three complexes but they may enhance the overall interaction between Arpp19 and the PP2A phosphatase. Noteworthy, the three conserved residues surrounding the phosphorylated serine, F60, D61 and D64 in human Arpp19, seems important for precise positioning of the serine into the catalytic site of PP2A. This nicely explained the impact of their mutation to alanine, that produce variants that are rapidly dephosphorylated by PP2A-B55 and lose their affinity for the phosphatase^16^.

**Figure 6.**
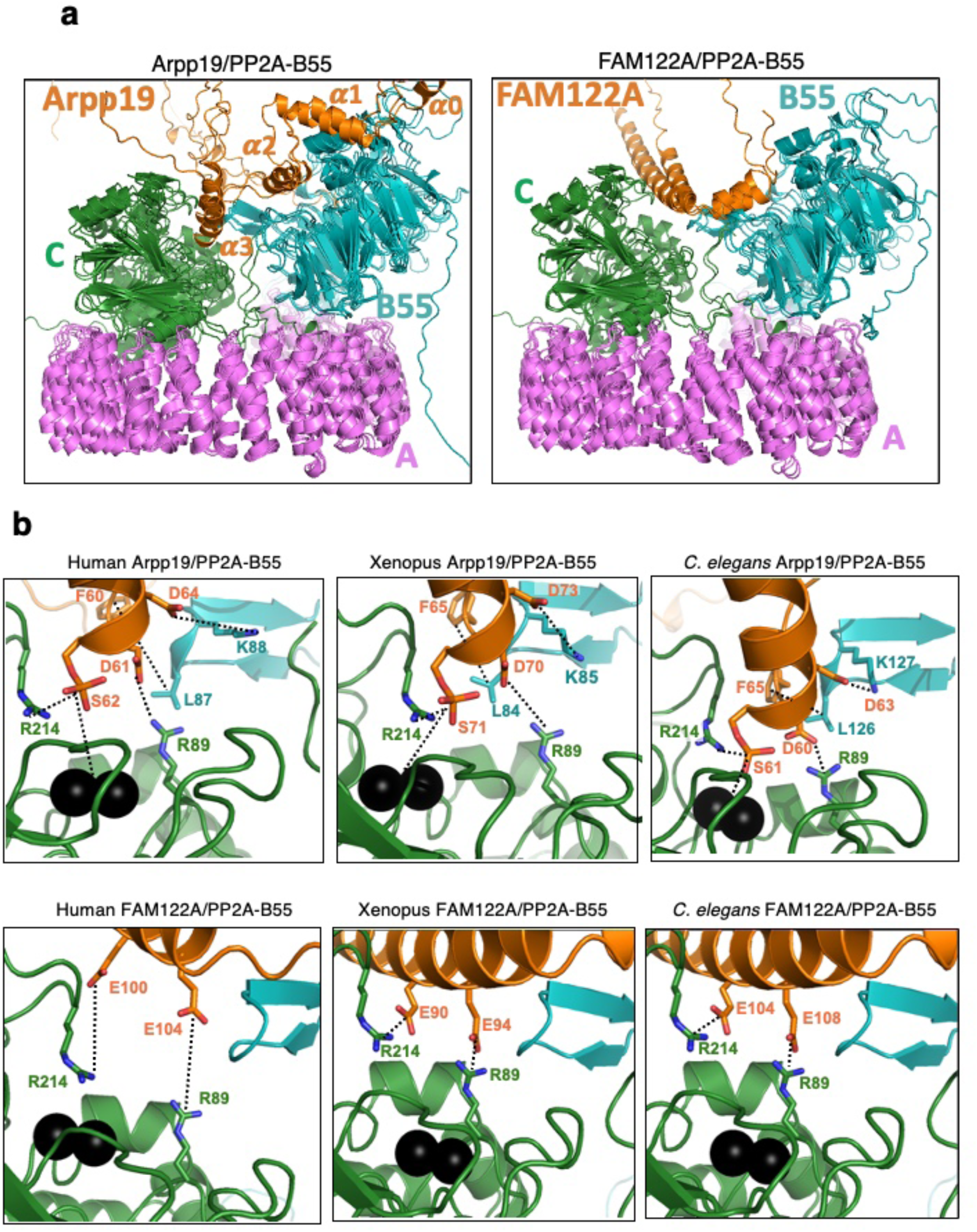
Arpp19 and FAM122A differently bind PP2A-B55. **(a)** Superposition of the models of Arpp19/ENSA and FAM122A in complex with ternary PP2A-B55 as predicted by Alphafold_multimer v 2.2 for three species (human, Xenopus and *C. elegans*). Arrp19/ENSA and truncated FAM122A are in orange ribbon while PP2A-B55 is in blue (B55), dark green (catalytic C subunit) and pink (the scaffold A subunit). **(b)** Zoom onto the entrance of the catalytic site of PP2A in complex with human Arpp19 and FAM122A. Manganese ions are shown as black spheres and sidechains are drawn as sticks with carbon colours as the corresponding polypeptide chains. Figure was drawn using Pymol (http://www.pymol.org).

Interestingly, no complexes could be predicted for the full-length FAM122A inhibitor with PP2A using Alphafold 2.2. On the contrary, short segments of FAM122A (from the three species used above) led to prediction of interactions between FAM122A and PP2A-B55. The interface partially overlaps with that observed for PP2A-Arpp19 complexes supporting the fact that the two inhibitors compete for PP2A-B55 binding. Nevertheless, the precise interactions differ significantly. First, FAM122A does not seem to point into the catalytic site as Arpp19 seems to do, although there are also two predicted salt bridges between two glutamate residues from FAM122A (E100 and E104 in human FAM122A) with the same arginines contacting D61 and S62 of Arpp19 of the catalytic subunit of PP2A (R89 and R214 in human PP2A) (Figure 6b). These two interactions are crucial since, as shown above (Figure 3f), the corresponding E90A and E94A Xenopus FAM122A mutants are unable to bind the phosphatase. Finally, as expected, the second helix of FAM122A containing the putative conserved SLiM motif, interacts with the B55 subunit and this interaction takes place in a region of B55 close to the one binding α2 helix of Arpp19.

In conclusion, our modeling data obtained for Arpp19 and FAM122A interaction with PP2A using Alphafold multimer suggest that the two inhibitors would produce analogous but distinct complexes with their common target. The size of these interfaces suggests that Arpp19 would be a stronger inhibitor than FAM122A and could thus promote the dissociation of the latter. This assumption is supported by newly predicted contacts with PP2A-B55 likely playing a major role in the strong inhibition of this enzyme by Arpp19 and that include the phosphorylation of the central serine that would tightly interact with the catalytic manganese ions.

### FAM122A is required *"in vivo"* for mitotic entry in the *C. elegans* germline

Our data above using ectopic FAM122A demonstrates that the addition of this protein to interphase Xenopus egg extracts promote mitotic entry. We thus sought to assess the impact of the depletion of the endogenous FAM122A from these extracts in mitotic entry. However, unfortunately, although we developed specific antibodies against this protein, they were unable to immunoprecipitate endogenous FAM122A. We thus used the nematode *Caenorhabditis elegans* as a model to investigate the role of FAM122A in PP2A-B55 inhibition and mitotic entry. The gene *F46H5.2* in *C. elegans* has been previously suggested as the putative orthologue of FAM122A protein^22^. To confirm that it does indeed correspond to *C. elegans* FAM122A (Ce FAM122A), we produced and purified this protein and we checked whether it was able to inhibit PP2A-B55 and to promote mitotic entry in interphase extracts. As shown in Figure 7a, dephosphorylation of S113 of Arpp19 and of T481 of PRC1 by PP2A-B55 was strongly delayed by the addition of this protein. Moreover, when supplemented to interphase extracts, Ce FAM122A induced mitotic entry and exit (Figure 7b). Thus, the product of the gene *F46H5.2* does correspond to the FAM122A orthologue.

**Figure 7.**
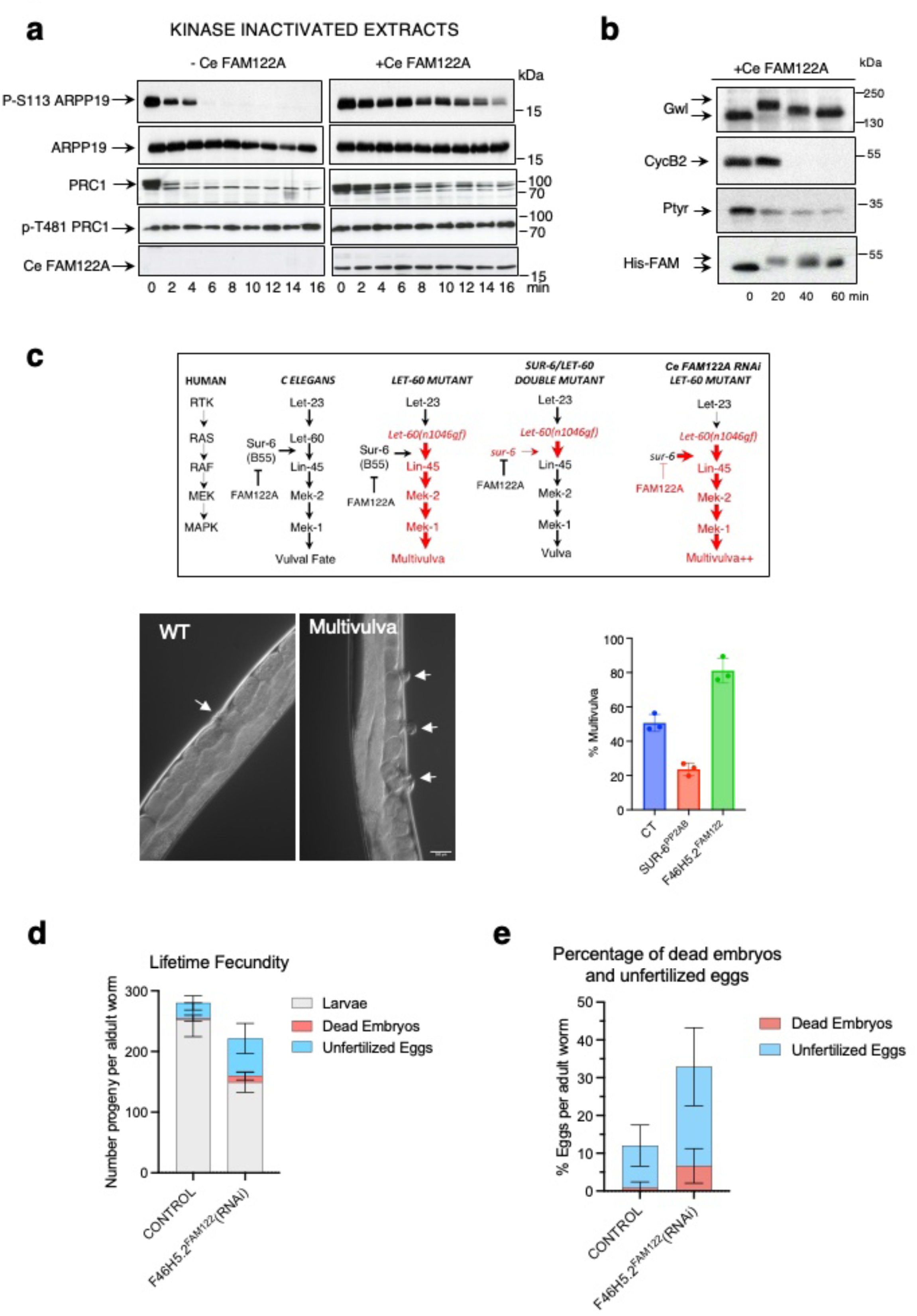
FAM122A is required to correctly enter into mitosis in *C elegans*. **(a)** Arpp19 or PRC1 *"in vitro"* phosphorylated by PKA or cdk1 respectively were supplemented together with Ce FAM122A to interphase extracts and the levels and dephosphorylation rate of S113 of Arpp19 or T481 of PRC1, as well as the amount of Ce FAM122A, were analysed by western blot. **(b)** Ce FAM122A was added to interphase extracts and the levels and phosphorylation of the indicated proteins measured by western blot. **(c)** Scheme representing the genetic dependences of the gain of function *let-60(gf)* (Ras) mutant and *sur-6* (B55 orthologue) mutant on multivulva phenotype in *C. elegans*. *sur-6* mutant (diminishes PP2A-B55 activity) on a *let-60(gf)* mutant background decreases the multivulva phenotype. Conversely, the depletion of *F46H5.2* (FAM122A orthologue) in a *let-60(gf)* mutant background significantly increases this phenotype suggesting that, as expected, *F46H5.2* protein would act as an inhibitor of PP2A-B55 in *C. elegans*. Big red arrows: increased multivulva phenotype. Shown are two representative images of a wildtype worm displaying one vulva or a mutant worm with a multivulva phenotype. The percentage of worms displaying multivulva phenotype in each mutant is represented as a mean+/- standard deviation. Significant differences calculated by non-parametric two tailed Mann-Whitney test are shown. **(d)** The number of larvae, dead embryos and unfertilized eggs were counted for control and *F46H5.2(RNAi)* worms and represented as mean values +/- standard deviation. **(e)** The percentage of dead embryos and unfertilized eggs were counted in worms at first day of adulthood and represented as for (d).

We then investigated the capacity of endogenous Ce FAM122A to inhibit PP2A-B55 "*in vivo"* in the worm. To this, we first took advantage of a multivulva phenotype induced by the mutant *let-60(n1046gf)*. This mutant, corresponding to a gain of function of the human orthologue Ras, promotes the formation of several vulvas in the worm (Figure 7c). It has been shown that the multivulva phenotype induced by *let-60(n1046gf)*^23^ can be attenuated by negatively modulating PP2A-B55 activity as this protein promotes Ras signalling during vulva development. Accordingly, we observed a suppression of the multivulva phenotype after depletion of SUR-6 (SUpressor of Ras-6, the orthologue of human PP2A-B55 subunit) by RNAi. In order to assess whether endogenous Ce FAM122A acts as an inhibitor of PP2A-B55, we examined the effect of the depletion of this protein by RNAi on the multivulva phenotype induced by the *let-60(n1046gf)* mutant. If this protein is a PP2A-B55 inhibitor *"in vivo"*, its depletion should promote the reactivation of this phosphatase and thus significantly increase the presence of multivulvas in the worms. Confirming this hypothesis, the depletion of Ce FAM122A by RNAi dramatically increased this phenotype (Figure 7c). Thus, Ce FAM122A does act as an inhibitor of PP2A-B55 "*in vivo"*.

We further explored the role of this protein in *C. elegans* in mitosis "*in vivo"*, in *F46H5.2^FAM^*^122^*^A^*(RNAi) treated worms. Interestingly these worms exhibit a decreased fecundity (Figure 7d) with a significant increased percentage of embryonic lethality at 25°C (median: 8,14% in RNAi-treated vs 0,30% in controls; p<0,0001) and a double number of unfertilized eggs compared to controls (average 26,18 % in RNAi-treated vs 12,10% in controls; p<0,014) (Figure 7e). Even wildtype *C. elegans* can produce unfertilized eggs when they age however, *F46H5.2^FAM^*^122^*^A^*(RNAi) worms display this phenotype on their first day of adulthood. These results suggest a fertility defect and led us to explore the function of the reproductive system. We thus investigated cell division within the *C. elegans* germline using nematode strains expressing histone and gamma-tubulin tagged with GFP under a germline and embryonic promoter. We focused on mitotic division of the germline stem cells of the distal gonad. Using chromatin and centrosome as proxy of mitotic progression^24^, we counted the number of stem cells in interphase and in mitosis (Supplementary Figure 3). We observed a significant reduction of interphase cells and an increase in mitotic cells in RNAi treated worms (Figure 8a). Interestingly, the number of cells in prophase (assessed by the presence of two separating centrosomes and a circular intact nuclei), was also dramatically increased, although some cells could progress into mitosis, probably due to a partial depletion of FAM122A. However, these cells displayed a perturbed mitotic progression since we observed a drop of the number of anaphases (Figure 8c). These data suggest that depletion of *F46H5.2* gene product decreases the capacity of germline stem cells to enter into mitosis and perturbs their progression through this phase of the cell cycle. To further investigate this hypothesis, we used a cell line with GFP-tagged-tubulin and with RFP-tagged histone in which we followed by time lapse microscopy the duration of mitosis. We considered mitosis duration as the time from first centrosome separation movement to anaphase onset. As depicted in Figure 8c, stem cells from control worms displayed a rapid mitosis, with a timing around 21 minutes. Conversely, Ce FAM122A depleted stem cells, stayed with separated centrosomes throughout the time of the experiment (40 minutes) indicating that these cells were arrested or highly delayed in prophase. Interestingly, as reported above, some cells could enter mitosis, however, although they did not display differences in the timing from nuclear envelope background to metaphase (congression), they showed a significant increase of the time required for anaphase onset. This delay is in agreement with the drop of the number of anaphase in RNAi-treated germline stem cells and suggests that FAM122A-dependent inhibition of PP2A-B55 could not be only required for mitotic entry, but could also be essential again during mitotic exit to promote anaphase onset when Gwl and cyclin B/cdk1 become inactivated by cyclin B proteolysis.

**Figure 8.**
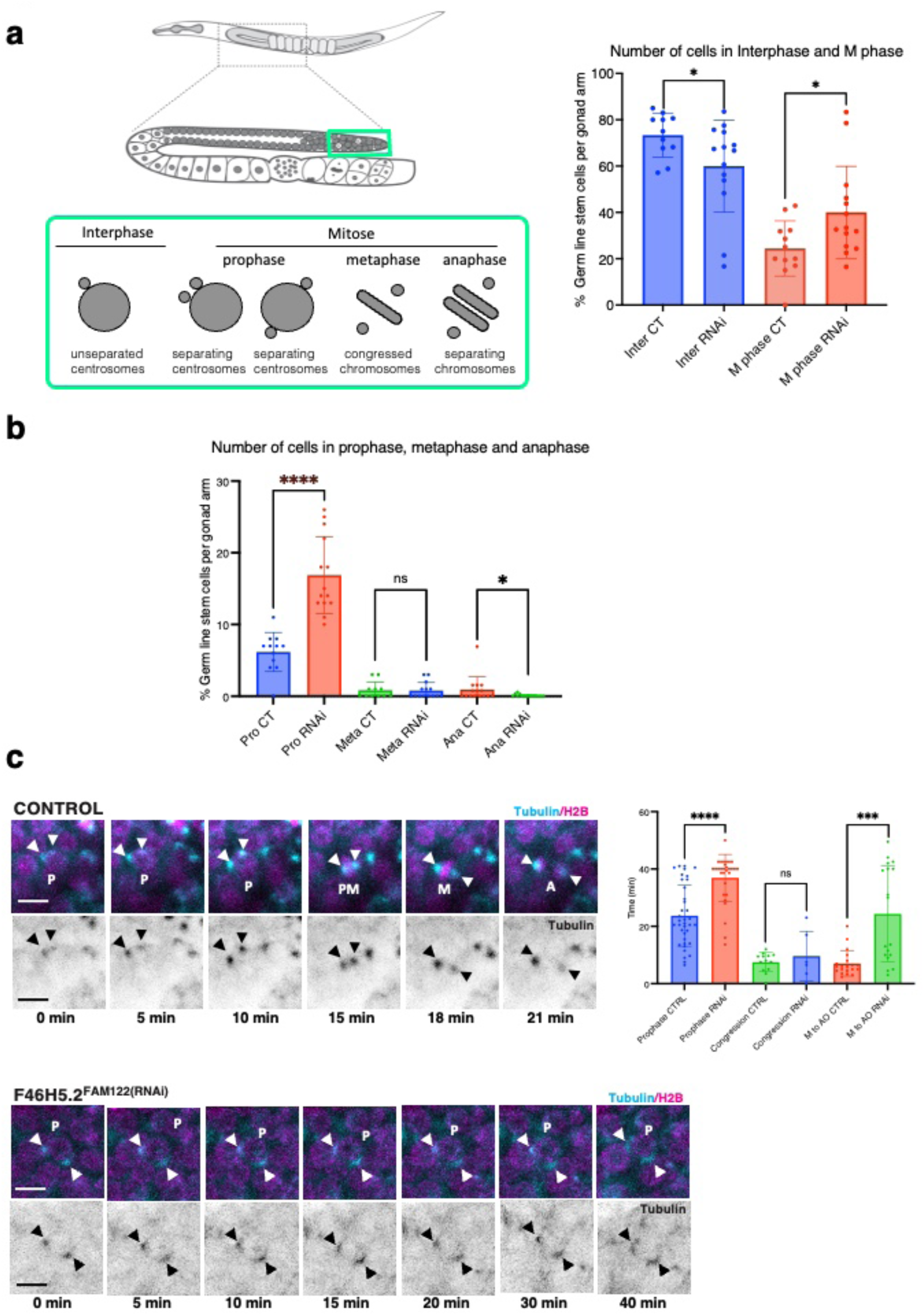
FAM122A is required "*in vivo"* in *C. elegans* to permit mitotic entry and progression in germ stem cells. **(a)** Scheme showing the position of germline stem cells within *C. elegans* gonad and the visualization of interphase, prophase, metaphase and anaphase cells using centrosomes and chromatin as the parameters to identify different phases of mitosis. Nematode strains expressing histone and gamma-tubulin tagged with GFP were immobilised and immediately imaged by confocal microscopy for the quantification of interphase and mitotic cells. Data are represented as mean +/- standard deviation. **(b)** Prophase, metaphase and anaphase germ stem cells in gonads of worms were counted as in (a) and represented as mean +/- standard deviation. **(c)** A GFP-tagged tubulin and with RFP-tagged histone nematode strain was followed by time lapse confocal microscopy and the timing of mitotic progression was determined in germ stem cells. Shown are representative images of stem cells performing mitotic division over time in control and RNAi treated worms. Timing of cells to perform prophase (from first centrosome movement to nuclear envelope breakdown), congression (from nuclear envelope breakdown to metaphase plate) or anaphase onset were recorded and represented as mean values +/- standard deviation. Significant differences calculated by non-parametric two tailed Mann-Whitney test are shown. Arrowheads: centrosomes. P:prophase. Sp: Spermatozoids. PM: prometaphase. A: anaphase. Scale bar: 10 μm.

## DISCUSSION

Protein phosphorylation plays a major role in the control of cell cycle progression. This phosphorylation results from a fine-tuned balance between the activities of cyclin/cdk complexes and phosphatases. The modulation of the phosphatase PP2A-B55 is key for S and M phases. During DNA replication, PP2A-B55 has to be inhibited to maintain the phosphorylation of the replication factor Treslin responsible of origin firing while at G2-M, the negative modulation of this phosphatase is essential for cyclin B/cdk1 activation.

To enter into mitosis a partial pool of cyclin B/cdk1 has to be turned on to induce, via Wee1/Myt1/Cdc25 phosphorylation, the amplification loop triggering cyclin B/cdk1 full activation. However, Wee1/Myt1/Cdc25 phosphorylation can be also achieved if PP2A-B55 is at least partially inhibited. This inhibition depends on the phosphorylation of Arpp19/ENSA and thus, requires the activity of its upstream kinase Gwl. However, at M entry, both Gwl and cyclin B/cdk1 are inactive and their activation depends on each other. Thus, none of them can be the initial event triggering mitosis. We hypothesize that this initial event corresponds to the accumulation of cyclin A/cdk1 activity. During G2, and in the presence of a FAM122A-dependent inhibition of PP2A-B55, cyclin A/cdk1 would be able to phosphorylate Cdc25 and Myt1 and to trigger cyclin B/cdk1 full activation and mitotic entry. Accordingly, because cyclin A/cdk1 escapes Myt1-dependent inhibition and FAM122A binding to PP2A-B55 does not require phosphorylation, their respective kinase and inhibitory activities precede Gwl and cyclin B/cdk1 activation. Supporting this idea, we show that the addition of FAM122A to interphase extracts promotes mitotic substrate phosphorylation, cyclin B/cdk1 activation and entry into mitosis. This phosphorylation is independent of cyclin B/cdk1 or Gwl activation but requires the presence of cyclin A/cdk1. We additionally show that FAM122A plays a key role in PP2A-B55 inhibition and mitotic entry "*in vivo*" in the *C. elegans* model in which, FAM122A depletion modulates phosphatase activity in vulval precursors and prevents mitotic entry in germ line stem cells.

However, strikingly, although FAM122A clearly binds PP2A-B55 during interphase, we discovered that this protein rapidly dissociated from the phosphatase as soon as the extract entered into mitosis and re-associated again upon mitotic exit. Our data show that this dissociation was induced by the phosphorylation of Arpp19 by Gwl. Accordingly, the addition of phospho-S71-Arpp19 to interphase extracts or to FAM122A-PP2A-B55 pull-downs promoted the exchange of FAM122A by phospho-S71-Arpp19 in the PP2A-B55 complex. These data indicate that, when phosphorylated by Gwl, Arpp19 competes with FAM122A and promotes its dissociation from the phosphatase. Indeed, our data support the hypothesis that phospho-S71-Arpp19 displays a much higher affinity, inhibitory activity and binding to PP2A-B55 than FAM122A. Accordingly, we estimated that, in our analysis, endogenous phospho-S71-Arpp19 is able to dissociate ectopic His-FAM122A from PP2A-B55 even when present in the extract at a molar concentration 187 times higher. These properties were supported by the structure predictions of FAM122A- and Arpp19-PP2A-B55 complexes obtained by AlphaFold2.2. Interestingly, structures predicted for FAM122-PP2A-B55 and phospho-Arpp19-PP2A-B55 complexes superimposed from three different species (human, Xenopus and *C. elegans*) revealed that FAM122A and phospho-S71-Arpp19 display different interactions with the PP2A-B55 heterocomplex. In this line, FAM122A weakly interacts with the C subunit via two glutamic acids forming salt bridges with two arginine residues of the C subunit of PP2A. These interactions locate the second alpha helix of this protein on the surface of the catalytic site occluding, in this way, substrate entry. Conversely, phospho-S71-Arpp19 tightly binds PP2A C mostly because of the presence of the phospho-serine residue that would directly interact with manganese ions of the catalytic site, an interaction that would be stabilized by two supplementary salt bridges formed with two arginine of the C subunit. This tight interaction can be only dissociated upon Arpp19 dephosphorylation, however, as we previously reported, dephosphorylation of the Arpp19 Gwl-site is very slow compared to regular substrates of this phosphatase making of this phosphoprotein a potent inhibitor of PP2A-B55^16^.

Although we do not know why FAM122A has to be dissociated from PP2A-B55 by phospho-Arpp19 (S71) during mitosis, we predict that this dissociation could be physiological relevant and required to induce a correct mitotic progression. In this line, it is possible that a higher PP2A-B55 inhibitory activity is required during mitotic progression to permit the late mitotic substrate phosphorylation. According with this hypothesis, previous data demonstrate that cyclin B1/cdk1 levels required to enter mitosis are lower than the amount of cyclin B1– cdk1 needed for mitotic progression^25^. Moreover, besides the differential inhibitory activity of phospho-S71-Arpp19, the fact that its dissociation from PP2A-B55 is gradually induced by dephosphorylation could be essential to stablish the temporal pattern of mitotic substrate dephosphorylation required for the correct order of mitotic events induced during mitotic exit.

In summary, we identified a new role of FAM122A in promoting G2-M transition through the inhibition of PP2A-B55 and we provided strong data indicating that cyclin A/cdk, together with FAM122A-dependent inhibition of PP2A-B55, are the initial events triggering mitotic entry. Finally, we showed that, upon mitotic entry, FAM122A is dissociated from PP2A-B55 by phospho-Arpp19 until the time when Gwl inactivation and Arpp19 becomes fully dephosphorylated permitting to FAM122A to re-associate again to the phosphatase.

## METHODS

### *“In vitro”* phosphorylation

Phosphorylation of Arpp19 on S71 or of Xenopus FAM122A by Gwl was induced by using GST-K72M hyperactive mutant form of Gwl purified from SF9 cells. For *“in vitro”* phosphorylation reaction, Xenopus His-Arpp19 protein or His-FAM122A and GST-K72M Gwl kinase were mixed at a final concentration of 2 μg/μl for the two first proteins and 45 ng/μl for the last one in a reaction buffer containing 7x10^-2^ μM ATPγ^33^P at a specific activity of 3000 Ci/ mmol, 200 μM ATP, 2 mM MgCl_2_ and Hepes 50 mM, pH 7,5.

For phosphorylation of Arpp19 on S113 by PKA, a final concentration of 0.5 μg/μl of Arpp19 and 50 ng/μl of His-PKA catalytic subunit purified from His-tag column were mixed to the reaction buffer containing 8x10^-3^ μM ATPγ^33^P at a specific activity of 3000 Ci/ mmol, 200 μM ATP, 2 mM MgCl_2_, 100 mM NaCl and 50 mM Tris pH 7.5.

For phosphorylation of GST-PRC1 on T481, cdk1 activity was obtained by immunoprecipitation. In brief, 1 μg of His-human cyclin A was mixed with 100 μl of CSF extracts during 30 minutes and subsequently supplemented with 50 μl of protein G magnetic Dynabeads pre-linked with 10 μg of Xenopus cdk1 C-terminus antibodies. After 45 minute-incubation the beads were washed 3 times with 500 mM NaCl, 50 mM Tris pH 7.5, twice with 100 mM NaCl, 50 mM Tris pH 7.5 and finally resuspended with 100 μl of reaction buffer containing 1mM ATP, 2 mM MgCl_2_, 100 mM NaCl and 50 mM Tris pH 7.5. GST-PRC1 was then added to the beads at a final concentration of 1,5 μg/ul.

For thio-phosphorylation of Arpp19 on S71, His-tag of His-Arpp19 protein was removed using TEV protease. Non-tagged Arpp19 was then used for phosphorylation at a final concentration of 1 μg/ μl in presence of 1mM ATPγS, 2 mM MgCl_2_, 100 mM NaCl, 50 mM Tris pH 7.5 and 45 ng/μl of GST-K72M Gwl kinase.

All *“in vitro”* phosphorylation reactions were incubated for 1 h, aliquoted and frozen at -70°C until use.

### Dephosphorylation reactions in kinase inactivated extracts

When the impact of Xenopus, human, or *C. elegans* FAM122A proteins on either S113 Arpp19 or T481 PRC1 dephosphorylation was checked in ATP-devoid interphase extracts, 1μl of the corresponding *“in vitro”* phosphorylation reaction was mixed to 1 μl (4.29 μg) of His-FAM122A, diluted with 9 μl of Tris 50 mM-10 mM EDTA buffer and supplemented with 10 μl of ATP-devoid interphase extracts adjusted to 500 mM NaCl with a solution of 5M NaCl and a sample of 2 μl was recovered at the indicated time-points. Time-point 0 min was prepared by adding separately 1 μl of extract and of 1 μl of phosphorylated substrate directly in Laemmli buffer.

### Immunoprecipitation/Immunodepletion

Immunodepletions were performed using 20 μl of extracts, 20 μl of G-magnetic Dynabeads (Life Technologies), and 2 μg of each antibody except for cyclin A for which we used 3,3 μl. Antibody-linked beads were washed 2 times with XB buffer, 2 times with Tris 50 mM, pH 7.5 and incubated for 15 min at room temperature (RT) with 20 μl of *Xenopus* egg extracts. The supernatant was recovered and used for subsequent experiments. To fully deplete endogenous proteins, two rounds of immunodepletion were performed.

For Gwl rescue experiments, 50 ng of K72M GWL were added to 20 μl of depleted egg extracts and a sample of 2 μl was recovered and used for western blot.

For histidine pulldown experiments, 20 μl of interphase or CSF extracts containing His-Xenopus, His-human or His-*C elegans* FAM122A proteins, were supplemented with 20 μl of HisPur^TM^NiNTA Magnetic beads (Life Technologies) and incubated for 10 minutes of continuous mixing at 21°C. Upon centrifugation, beads are washed twice with XB buffer and supplemented with Laemmli sample buffer for western blot use.

### Protein purification

His-Xenopus Arpp19, His-Xenopus wildtype and mutant of FAM122A in which all serine and threonine residues were mutated into alanine, as well as, His-human FAM122A, His-*C elegans* FAM122A, His-human PRC1, and His-Rat Catalytic Subunit of PKA were produced in *Escherichia coli* and purified using TALON Superflow Metal Affinity Resin. GST-Xenopus FAM122A protein was produced in *Escherichia coli* and purified using a glutathione column.

### CSF, interphase and kinase-inactivated egg extracts

Frogs were obtained from « TEFOR Paris-Saclay, CNRS UMS2010 / INRAE UMS1451, Université Paris-Saclay», France and kept in a *Xenopus* research facility at the CRBM (Facility Centre approved by the French Government. Approval n° C3417239). Females were injected of 500 U Chorulon (Human Chorionic Gonadotrophin) and 18 h later laid oocytes were used for experiments. Adult females were exclusively used to obtain eggs. All procedures were approved by the Direction Generale de la Recherche et Innovation, Ministère de L’Enseignement Supérieur de la l’Innovation of France (Approval n° APAFIS#40182-202301031124273v4).

Metaphase II-arrested egg extracts (CSF extracts) were obtained by crushing metaphase II-arrested oocytes in the presence of EGTA at a final concentration of 5mM^26^. Interphase Xenopus egg extracts obtained from metaphase II-arrested oocytes 35 minutes after Ca^2+^ Ionophore (final concentration 2 μg/ml) treatment.

Kinase-inactivated egg extracts were obtained from dejellied unfertilised eggs transferred into MMR solution (25 mM NaCl, 0.5 mM KCl, 0.25 MgCl_2_, 0.025 mM NaEGTA, 1.25 mM HEPES-NaOH pH7.7), washed twice with XB Buffer (50 mM sucrose, 0.1 mM CaCl_2_, 1 mM MgCl_2_, 100 mM KCl, HEPES pH 7.8) and centrifuged twice for 20 minutes at 10 000 g. Cytoplasmic fractions were then recovered, supplemented with RNAse (10 μg/ml final concentration) and dialyzed versus a solution of 50 mM Tris pH 7.7, 100 mM NaCl overnight to eliminate ATP. Upon dialysis, extracts were ultracentrifuged for 50’ at 300 000 g and supernatant recovered for use.

### Plasmids

Xenopus wildtype and the Δ(1-73) and serine/threonine-to-alanine mutant form (accession number NP_001085566.1) cDNAs, as well as human (accession number NP_612206.5), and *C. elegans* (accession number NP_001024675.1) cDNAs were synthesized by GeneCust (France) and subcloned into the HindIII-XhoI site of pET14b for human FAM122A, into the NdeI-BamHI site of pET14b for *C elegans*, into the XhoI-BamHI site of pET14b for Xenopus wildtype and mutant forms and into the BamHI-XhoI site of pGEX4T1 for wildtype Xenopus FAM122A.

### Antibodies

Xenopus, human and *C elegans* FAM122A protein was detected using anti-histidine antibodies. Antibodies used in this study are detailed in Supplementary material, Table 1.

### Mutagenesis

Deletions and single-point mutations of Xenopus FAM122A were performed using Pfu ultra II fusion DNA polymerase. Oligonucleotides were purchased from Eurogentec and are detailed in the Supplementary material, Table 2.

### *C. elegans* culture and RNAi mediated depletion

*C. elegans* worm strains N2 (wildtype ancestral, Bristol) and MT2124 (*let-60(gf)*) were obtained from the CGC (https://cgc.umn.edu) and maintained at 20 °C on NGM plate using standard procedures^27^ except that worms were fed with HT115 bacteria to standardize their growth condition with the RNAi mediated depletion. RNAi feeding was performed as described previously ^28^. HT115 thermocompetent *E. coli* were transformed with L4440 empty vector (control) or containing a sequence targeting, SUR-6^PP2AB^ or F46H5.2^FAM122^. The L4440 vector targeting F46H5.2^FAM122^ was generated using a fragment amplified from *C. elegans* cDNA corresponding to 715 bp of the coding sequence (nucleotides 398 to 1112) which was subsequently inserted between XhoI and NotI restriction enzyme sites in L4440 vector. In addition, a vector targeting PLK-1 was used to control for RNAi efficiency as depletion of this protein during larval development induce 100 % protruded vulva and sterility. Control, SUR-6^PP2AB^ and PLK-1 vectors come from the Arhinger’s library. In a 14 ml culture tube, 2 ml of LB medium supplemented with 100 µg/mL of ampicillin were inoculated with a colony of HT115 bacteria transformed with respective L4440 vectors and incubated at 37 °C under agitation. After 7 h, 200 µl of this bacterial culture was transferred on a 60 mm diameter NGM plate containing 0.2 mM IPTG and 50 µg/ml of carbenicillin. Plates were allowed to dry overnight at room temperature, stored at 4 °C and used within 48 h. Worms were synchronized using the alkaline bleach method (1.2% NaOCl, 250 mM KOH in water^29^). Eggs obtained from the alkaline bleach were allowed to hatch overnight at 16 °C in M9 buffer. To score FAM122A effect on Ras (*let-60(gf))* induced multivulva phenotype, around 200 synchronized larvae were placed on a single feeding plate and incubated 66-72 h at 20 °C until control worms reach adulthood. Phenotypes were scored at the time where the first eggs laid by control worms started to hatch to ensure that all worms fully develop to the adult stage. Scoring was only performed if 100% protruded vulva and 100% sterility was obtained on PLK-1 plates. Vulval defect phenotypes was scored under a dissecting scope by counting the total number of worms and the number of worms exhibiting multiple vulva. The counting was repeated once for each plate to minimize scoring errors. Images representative of vulval phenotypes were acquired with a scMOS ZYLA 4.2 M camera on a Zeiss Axioimager Z2 and a 20X Plan Apochromat 0.8 NA using worms immobilized with a solution of 10 mM sodium azide and mounted on an 3 % agarose pad in between a microscopy slide and a cover glass.

### Live imaging of *C. elegans*

For "*in vivo"* imaging, we used nematode strains expressing GFP-gamma-tubulin and GFP-histone (TH32: unc-119(ed3), ruIs32[pAZ132; pie-1/GFP::histone H2B] III; ddIs6[GFP::tbg-1; unc-119(+)] V or GFP-tubulin and histone-mCherry (JDU19: *ijmSi7 [pJD348; Pmex-5_gfp::tbb-2; mCherry::his-11; cb-unc-119(+)] I; unc-119(ed3) III)* and JDU233: *ijmSi63 [pJD520; mosII_5’mex-5_GFP::tba-2; mCherry::his-11; cb-unc-119(+)] II; unc-119(ed3) III*) kindly provided by Julien Dumont (Institut Jacques Monod, Paris). Worms were cultured at 25 °C and bacterial feeding was performed similarly to the multivulva experiment, except that worms were not synchronized by alkaline bleach. Instead, 6 worms were allowed to lay eggs on a 60 mm diameter NGM feeding plate. After 2 h at 25 °C adults were removed and their progeny was allowed to develop on the feeding plate at 25 °C for 44-52 h until the first adults start to lay eggs (young adults). Worms were anesthetized in 0.02 % tetramisole in M9 buffer for 10 min before being transferred onto a 4 % agarose pad ^30^. The pad was then covered with 22x22 mm coverslip and sealed with VaLaP (vaseline and paraffin wax and lanolin, 1:1:1). The chamber was filled with M9 to prevent drying and to dilute tetramisole. Immobilized worms were imaged for a maximal time of 45 min and were not maintained more than an hour after being removed from their feeding plate. Germline stem cell division imaging was performed according to Gerhold et al.^24^. To minimize toxicity, we used minimal laser intensity and exposure using a spinning disk confocal Yokogawa W1 on an Olympus inverted microscope coupled to a sCMOS Fusion BT Hamamatsu camera. Long-term live imaging (up to 45 min) was performed with a 30x UPLSAPO 1.05 NA DT 0.73 mm silicone objective by taking a full z-stack of the entire worms (∼40 µm) with 2 µm steps every 20 s or 30 s. For higher resolution images and to score mitotic phases using centrosome and chromatin markers, a single z-stack was performed on each worm with a 1 µm z-section and using a 60x UPLSAPO 1.3 NA DT 0.3mm silicone objective.

### Analysis of germline stem cells divisions

Images and movies were visualized using ImageJ or CellSens visualization software (OlyVIA, Olympus). Analysis of germline stem cells divisions was made following a previous article from Gerhold et al. ^24^. Accordingly, timing of prophase corresponds to the initiation of centrosome separation (GFP-tubulin) to the beginning of chromosome congression monitored using the mCherry-histone. As ovulation generates movement within the gonad arm, cells can move in 3D, therefore tracking of individual cell progression was performed by navigating throughout z-stack over time. Representative images and montages in the Figure 8 correspond to a maximal intensity projection of 4-5 z-sections (8-10 µm). To quantify the number of cells in each respective cell cycle phases, we used GFP-histone and GFP-gamma-tubulin signal to monitor centrosome separation, nuclear envelope breakdown, chromosome congression and separation. The number of stem cells in the mitotic zone of the distal gonad was estimated by counting the number of nuclei based on histone signal. Here, cells were considered in prophase when having 2 separating or opposed centrosomes (spots) closed to nuclei or condensed chromosomes. Metaphase and anaphase correspond respectively to a single plate or two separated plates of tightly condensed chromosomes.

### Protein structure modelling

Complexes of PP2A-B55 with Arpp19/ENSA or FAM122A were produced using Alphafold_Multimer version 2.2 (https://www.biorxiv.org/content/early/2021/10/04/2021.10.04.463034) with the protein sequences from three species (*C. elegans, X. laevis* and *H. sapiens*). Manganese ions were added to the resulting models by superposition and extraction from the crystal structure of PP2A (PDB3DW8). The phosphoserine was modelled by superposing a phosphoserine onto the corresponding serine in each complex of interest and searching for the best orientation using our recent webserver ISPTM2 (https://isptm2.cbs.cnrs.fr). Because full-length FAM122A sequences led to no interactions as predicted by Alphafold 2.2, truncated version of FAM122A were used instead. For the human, Xenopus, and worm sequences, the shortened versions comprised residues 84 to 117, 61 to 120 and 70 to 149, respectively.

## DATA AVAILABILITY

All data that support the findings of this study are provided as a Source Data file and are available from the corresponding authors upon a reasonable request. There are no restrictions on data availability. Source data are provided with the paper.

Accession Codes are availables for Xenopus FAM 122A, Human FAM 122A and *C.elegans* FAM 122A at:

Xenopus FAM122A : PABIR family member 3 L homeolog

https://www.ncbi.nlm.nih.gov/nucleotide/NM_001092097.1?report=genbank&log$=nuclalign&blast_rank=1&RID=83FAFZZ0013

Human Fam122A : PPP2R1A-PPP2R2A-interacting phosphatase regulator 1

https://www.ncbi.nlm.nih.gov/protein/NP_612206.5?report=genbank&log$=protalign&blast_rank=1&RID=83FJ2J02016

C.elegans FAM122A F46H5.2, isoform a https://www.ncbi.nlm.nih.gov/protein/NP_001379960.1?report=genbank&log$=protalign&blast_rank=1&RID=83G2ABGU013

## ACKNOWLEDGMENTS

We are grateful to Marc Plays and Phillipe Richard for animal and antibody production and to the imaging facility MRI, member of the France-BioImaging national infrastructure supported by the French National Research Agency (ANR-10-INBS-04, «Investments for the future»). Some nematode -strains were provided by the CGC, which is funded by NIH Office of Research Infrastructure Programs (P40 OD010440). We thank Julien Dumont (Institut Jacques Monod, Paris) for kindly providing GFP-gamma-tubulin and mCherry-histone nematode strains and Lucie Van Hove, Sylvain Roque, Morgane Robert and Celia Benchoug for their technical help. This work was supported by the Agence National de la Recherche (REPLIGREAT, ANR-18-CE13-0018-01, MITODISSECT, ANR-22-0022 and MTDiSco, ANR-20-CE13-0033), La Ligue Nationale Contre le Cancer (Equipe Labellisée, EL2019 CASTRO and EL2018 PINTARD), Ligue Nationale Contre le Cancer (Comité Département 66/ LACROIX), the French Infrastructure for Integrated Structural Biology (FRISBI, ANR-10-INSB-005 and the infrastructure ChemBioFrance.

## AUTHOR CONTRIBUTION

B.L designed and performed most of the *C. elegans* experiments and analysed data. LP designed and performed some others *C. elegans* experiments and analysed data. S.V., designed and performed biochemical experiments in interphase and CSF extracts. J.C.L., designed and performed all dephosphorylation assays. Lucie Van Hove (LP lab) conceived and provided the RNAi feeding vector targeting FAM122A. G.L., performed modelling and wrote modelling section. A.C. designed some experiments, analysed data and wrote the paper. T.L. designed the experiments, analysed data and wrote the paper.

## SUPPLEMENTARY FIGURES

**Supplementary Figure 1:**
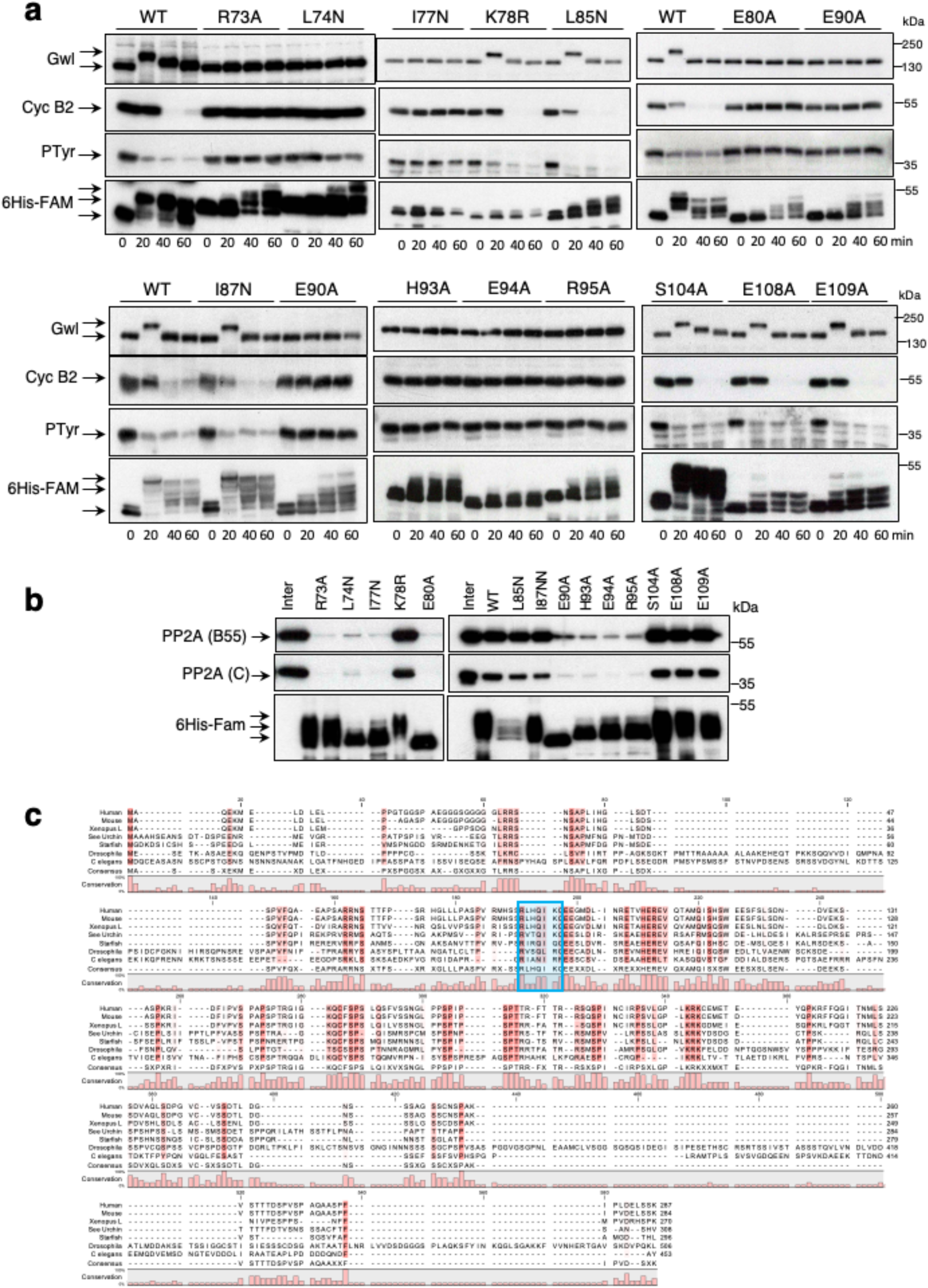
Identification of the residues of αH1 and αH2 required to PP2A inhibitory activity of Xe FAM122A. **(a)** The capacity of the indicated mutants of Xe FAM122A to promote mitotic entry when supplemented to interphase extracts is shown. **(b)** B55/C binding to the single mutants of the αH1 and αH2 is measured by His-pull down. **(c)** FAM122A sequence alignment on different species. Putative SLiM motif is highlighted in blue.

**Supplementary Figure 2:**
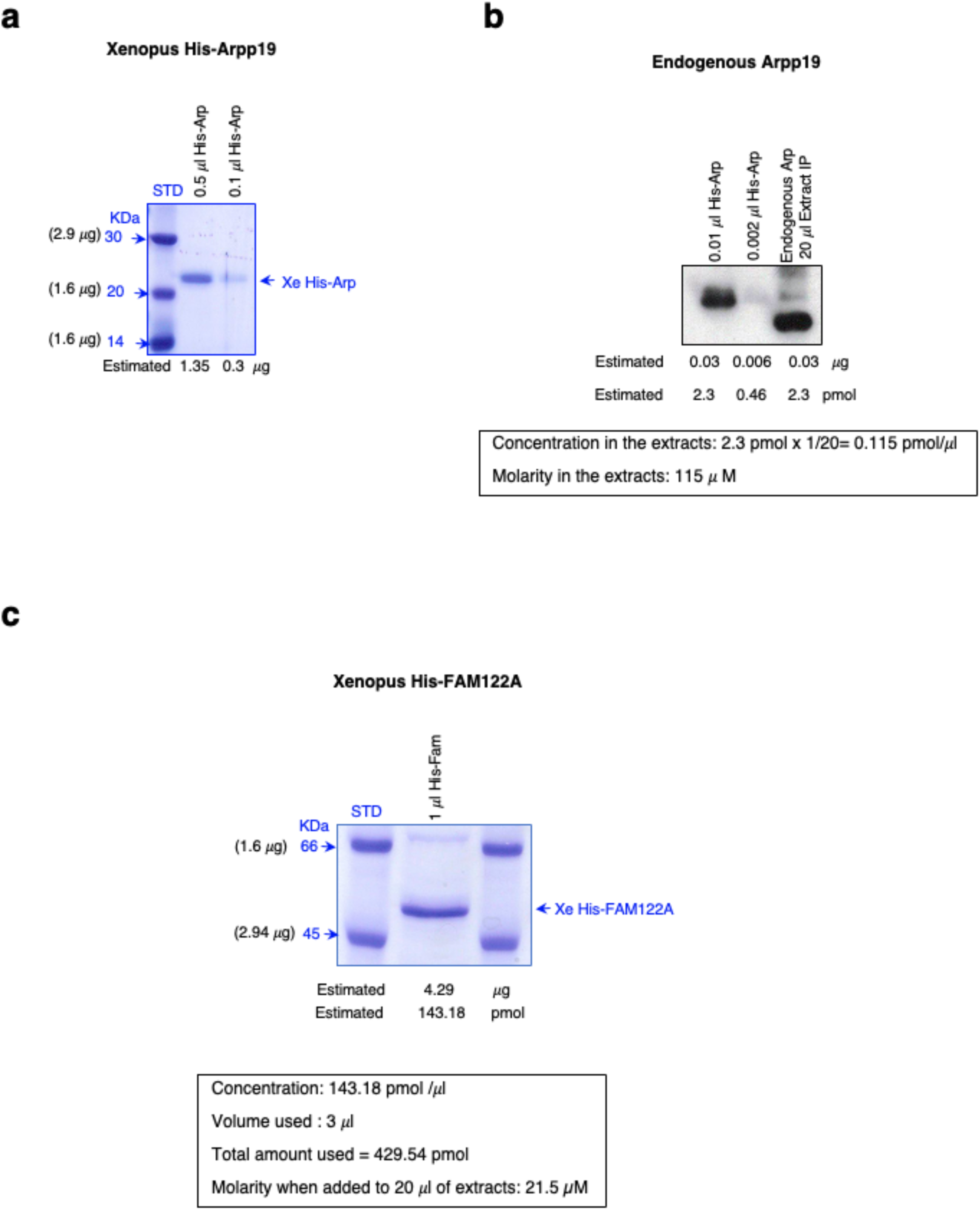
Estimation of final concentration in the extracts of endogenous Arpp19 and of the His-FAM122A used to induce mitotic entry. **(a)** The indicated volumes of the His-Arpp19 purified protein were used for PAGE-SDS and Coomassie blue staining. Intensity bands corresponding to His-Arpp19 were compared to standard markers and the protein amount quantified using ImageJ and estimated. **(b)** 20 μl of extracts were immunoprecipitated using anti-Arpp19 antibodies and subjected to western blot and the Arpp19 signal was compared with the one obtained for 0,03 and 0,006 μg of recombinant Xe-His-Arpp19. **(c)** Xenopus His-FAM122A concentration was estimated as for (a) and the final molarity corresponding to 3μl of His-FAM122A in 20 μl of extracts used for the experiments calculated.

**Supplementary Figure 3:**
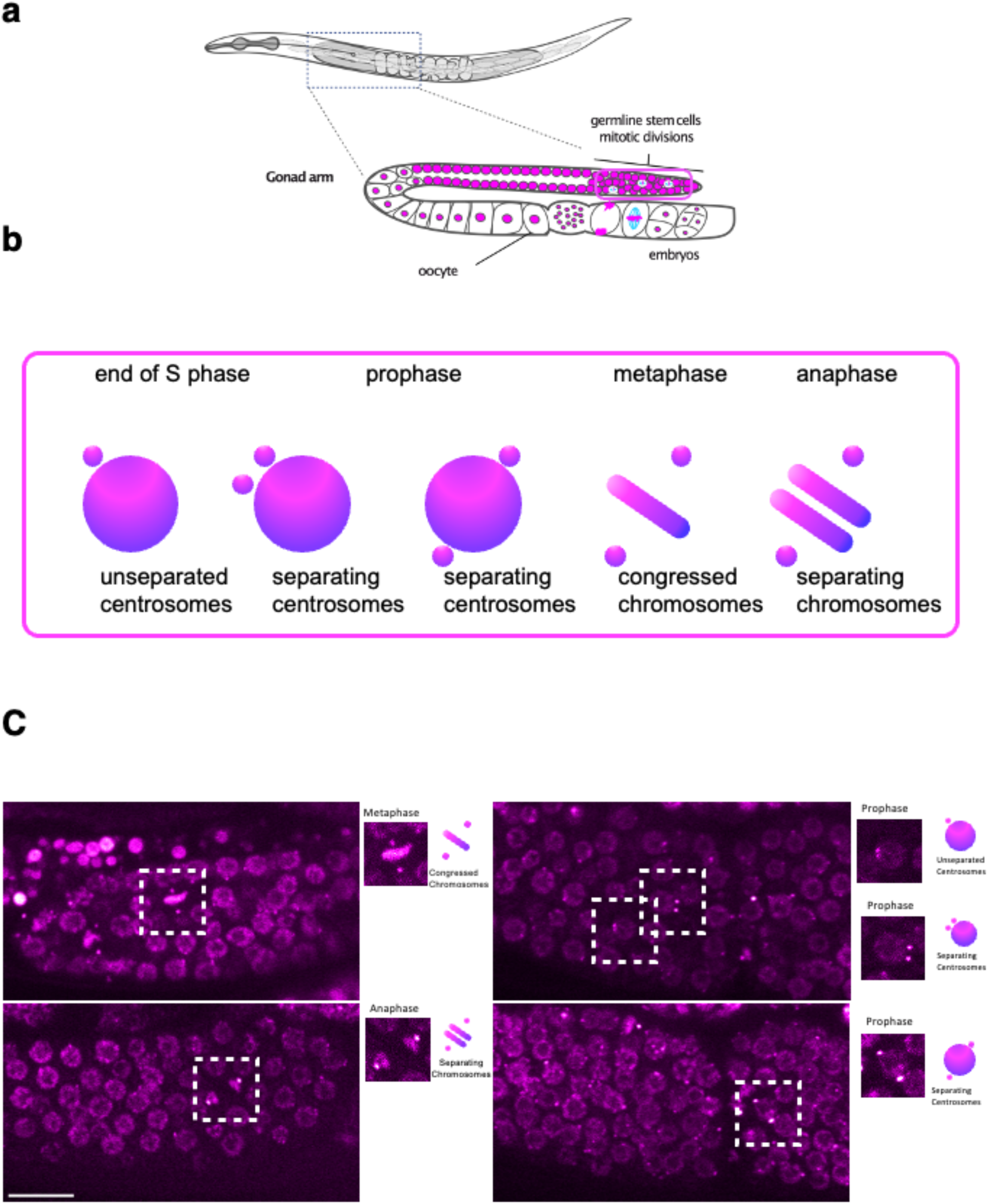
CeFAM122A depletion impairs mitotic entry of germ line stem cells in *C. elegans* nematodes. **(a)** Schematic representation of *C. elegans* nematode reproductive system. Germ line stem cells are located in the most distal part of each gonad arm. Chromatin is represented in magenta, microtubules in cyan **(b)** Schematic representation of the mitotic features observed and quantified based on chromatin and gamma-tubulin markers in Figure 8. **(c)** Representative images of the germ line stem cells imaged in TH32 strain (GFP-gamma-tubulin and GFP-histone) in Ce FAM122 depleted worms (right panel) or control worms (left panel). White dashed squares represent representative mitotic phases highlighted in zoomed square images next to each panel.

## REFERENCES

1. Mochida, S., Maslen, S. L., Skehel, M. & Hunt, T. Greatwall Phosphorylates an Inhibitor of Protein Phosphatase 2A That Is Essential for Mitosis. Science 330, 1670–1673 (2010).

2. Gharbi-Ayachi, A. et al. The Substrate of Greatwall Kinase, Arpp19, Controls Mitosis by Inhibiting Protein Phosphatase 2A. Science 330, 1673–1677 (2010).

3. Mochida, S., Ikeo, S., Gannon, J. & Hunt, T. Regulated activity of PP2A-B55 delta is crucial for controlling entry into and exit from mitosis in Xenopus egg extracts. The EMBO journal 28, 2777–85 (2009).

4. Vigneron, S. et al. Greatwall maintains mitosis through regulation of PP2A. EMBO J 28, 2786–93 (2009).

5. Pomerening, J. R., Sontag, E. D. & Ferrell, J. E. Building a cell cycle oscillator: hysteresis and bistability in the activation of Cdc2. Nat. Cell Biol. 5, 346–351 (2003).

6. Sha, W. et al. Hysteresis drives cell-cycle transitions in Xenopus laevis egg extracts. Proceedings of the National Academy of Sciences 100, 975–980 (2003).

7. Burgess, A. et al. Loss of human Greatwall results in G2 arrest and multiple mitotic defects due to deregulation of the cyclin B-Cdc2/PP2A balance. Proceedings of the National Academy of Sciences of the United States of America 107, 12564–9 (2010).

8. Hached, K. et al. ENSA and ARPP19 differentially control cell cycle progression and development. The Journal of Cell Biology 218, 541–558 (2019).

9. Vigneron, S. et al. Characterization of the Mechanisms Controlling Greatwall Activity. Molecular and Cellular Biology 31, 2262–2275 (2011).

10. Blake-Hodek, K. A. et al. Determinants for activation of the atypical AGC kinase Greatwall during M phase entry. Molecular and cellular biology 32, 1337–53 (2012).

11. Alvarez-Fernandez, M. et al. Greatwall is essential to prevent mitotic collapse after nuclear envelope breakdown in mammals. Proceedings of the National Academy of Sciences of the United States of America 110, 17374–9 (2013).

12. Coulonval, K., Bockstaele, L., Paternot, S. & Roger, P. P. Phosphorylations of cyclin-dependent kinase 2 revisited using two-dimensional gel electrophoresis. J Biol Chem 278, 52052–52060 (2003).

13. Booher, R. N., Holman, P. S. & Fattaey, A. Human Myt1 Is a Cell Cycle-regulated Kinase That Inhibits Cdc2 but Not Cdk2 Activity*. Journal of Biological Chemistry 272, 22300–22306 (1997).

14. Fan, L. et al. FAM122A, a new endogenous inhibitor of protein phosphatase 2A. Oncotarget 7, 63887–63900 (2016).

15. Li, F. et al. CHK1 Inhibitor Blocks Phosphorylation of FAM122A and Promotes Replication Stress. Mol Cell 80, 410–422.e6 (2020).

16. Labbé, J. C. et al. The study of the determinants controlling Arpp19 phosphatase-inhibitory activity reveals an Arpp19/PP2A-B55 feedback loop. Nat Commun 12, 3565 (2021).

17. Lemonnier, T., et al. The M-phase regulatory phosphatase PP2A-B55δ opposes protein kinase A on Arpp19 to initiate meiotic division. http://biorxiv.org/lookup/doi/10.1101/810549 (2019) doi:10.1101/810549.

18. Cundell, M. J. et al. The BEG (PP2A-B55/ENSA/Greatwall) Pathway Ensures Cytokinesis follows Chromosome Separation. Molecular Cell 52, 393–405 (2013).

19. Vigneron, S. et al. Cyclin A-cdk1-Dependent Phosphorylation of Bora Is the Triggering Factor Promoting Mitotic Entry. Dev. Cell 45, 637–650.e7 (2018).

20. Fowle, H. et al. PP2A/B55α substrate recruitment as defined by the retinoblastoma-related protein p107. eLife 10, e63181 (2021).

21. Williams, B. C. et al. Greatwall-phosphorylated Endosulfine is both an inhibitor and a substrate of PP2A-B55 heterotrimers. eLife 3, e01695 (2014).

22. Kim, W., Underwood, R. S., Greenwald, I. & Shaye, D. D. OrthoList 2: A New Comparative Genomic Analysis of Human and Caenorhabditis elegans Genes. Genetics 210, 445–461 (2018).

23. Sieburth, D. S., Sundaram, M., Howard, R. M. & Han, M. A PP2A regulatory subunit positively regulates Ras-mediated signaling during Caenorhabditis elegans vulval induction. Genes Dev 13, 2562–2569 (1999).

24. Gerhold, A. R., et al. Investigating the Regulation of Stem and Progenitor Cell Mitotic Progression by In Situ Imaging. Current Biology 25, 1123–1134 (2015).

25. Lindqvist, A., Zon, W. van, Rosenthal, C. K. & Wolthuis, R. M. F. Cyclin B1–Cdk1 Activation Continues after Centrosome Separation to Control Mitotic Progression. PLOS Biology 5, e123 (2007).

26. Lorca, T. et al. Constant regulation of both the MPF amplification loop and the Greatwall-PP2A pathway is required for metaphase II arrest and correct entry into the first embryonic cell cycle. Journal of cell science 123, 2281–91 (2010).

27. Brenner, S. The genetics of Caenorhabditis elegans. Genetics 77, 71–94 (1974).

28. Kamath, R. S., Martinez-Campos, M., Zipperlen, P., Fraser, A. G. & Ahringer, J. Effectiveness of specific RNA-mediated interference through ingested double-stranded RNA in Caenorhabditis elegans. Genome Biol 2, RESEARCH0002 (2001).

29. Stiernagle, T. Maintenance of C. elegans. WormBook 1–11 (2006) doi:10.1895/wormbook.1.101.1.

30. Laband, K., Lacroix, B., Edwards, F., Canman, J. C. & Dumont, J. Live imaging of C. elegans oocytes and early embryos. Methods Cell Biol 145, 217–236 (2018).

